# Change in liana density over 30 years in a Bornean rain forest supports the escape hypothesis

**DOI:** 10.1101/2020.08.03.234500

**Authors:** D. M. Newbery, C. Zahnd

## Abstract

Liana dynamics may influence tree dynamics and vice versa. Only long-term studies can perhaps disentangle them. In two permanent plots of lowland dipterocarp forest at Danum a liana census in 1988 was repeated in 2018. The primary forest was still in a late stage of recovery from an inferred large and natural disturbance in the past. Mean number of lianas per tree decreased by 22 and 34% in plots 1 and 2, and in different ways. By 2018 there were relatively more trees with few lianas and relatively fewer trees with many lianas than in 1988. The redistribution was strongest for overstorey trees of the Dipterocarpaceae (more with no lianas by 2018) and understorey trees of the Euphorbiaceae (many losing high loads in especially plot 2). Proportion of trees with lianas increased overall by 3.5%. Number of lianas per tree showed a quadratic relationship with tree size: maximal for large trees, fewer for smaller and very large trees. Tree survival and stem growth rate were significantly negatively related to the number of lianas after accounting for spatial autocorrelation. Monte Carlo random subsampling of trees in 1988 and 2018, to achieve statistical independence, established significance of change. Dipterocarps and euphorbs clearly differed in their liana dynamics between plots. Regression models had different forms for the two plots, which reflected a complicated structural-spatial variability in host-liana dynamics. Analysis of the abundant tree species individually, highlighted a group of emergent dipterocarps with low liana counts decreasing with time. Building on an earlier hypothesis for this forest type and site, these very large trees appear to have been losing their lianas by branch shedding, as they moved into and out of the main canopy. They were evidently ‘escaping’ from the parasite. The process may in part explain the characteristically very uneven forest canopy at Danum. Change in liana density was therefore contingent on both forest history and site succession, and on plot-level structure and tree dynamics. Liana promotion in the intermittent ENSO dry periods was seemingly being offset by closing of the forest and dominance by dipterocarps in late seral stages.

## INTRODUCTION

Lianas, or woody vines (climbers), are a conspicuous component of most tropical forests (Richards 1996). They are structural parasites: climbing trees allows them to reach the forest canopy with minimal investment in mechanical support. In the canopy they compete with trees for light (Schnitzer and Bongers 2002) and thereby affect tree growth, survival, recruitment and reproduction negatively (Putz 1984, Clark & Clark 1990, Phillips et al. 2005, van der Heijden & Phillips 2009, Ingwell *et al*. 2010). Susceptibility and tolerance to liana infestation can vary between tree species, leading to changes in forest composition, structure and population dynamics (Hegarty 1991, Campbell & Newbery 1993, Visser *et al*. 2018b, Muller-Landau and Visser 2019). Lianas fit well into a broader host-parasite theoretical modelling framework (Muller-Landau and Pacala 2020).

Several studies have shown increased liana abundance in rainforests across Central and Southern America in recent decades. Phillips et al. (2002) and Wright et al. (2004) showed that abundances of lianas increased significantly in E. Amazonia and Panama, respectively. Six more studies have broadly confirmed this trend in the neotropics (Schnitzer and Bongers 2002). In the paleotropics, three studies in Africa all showed decreased liana infestation (Caballé & Martin 2001, Thomas *et al*. 2015, Bongers et al. 2020), as did the one only study in SE Asia (Wright *et al*. 2015). These four paleotropical cases though, as with the ones from the neotropics, vary so widely in their site conditions, historically contingent factors, sampling approaches, and ways of recording lianas abundance, that drawing generalizations is at present unreliable. African rain forests, for instance, like the C. and S. American ones, experience one sharp or, more often, two moderate dry seasons per year, those in SE Asia very slight or no seasonality (Walsh 1996a).

Liana densities are most prominent in seasonal tropical forests (Gentry 1991, DeWalt et al. 2015), occur extensively in secondary successional stages of forest recovery (DeWalt et al. 2000, Wright 2005a,), but also increase in large tree fall gaps in late successional forest (Whitmore 1984, Schnitzer and Carson 2010). Three climate-related processes or factors have been proposed to explain liana increases in neotropical, primary forests (i) Increased rates of natural perturbation and disturbance due to droughts and storms, which open up canopies and create more gaps; (ii) Increased evapotranspiration due to longer, more intense, dry seasons, which favour lianas because of their deep rooting, stem anatomical adaptations, and highly efficient use of water, and (iii) increased global atmospheric carbon dioxide levels, which promotes the growth of lianas faster than it does of trees, especially under more lighted conditions (Schnitzer 2005, Swaine and Grace 2007, Cai et al. 2009, Schnitzer and Bongers 2011).

Lianas are nevertheless an important feature of aseasonal and less-recently disturbed rain forests in the paleotropics, particularly in SE Asia where primary forests are composed in recent decades largely of late-successional stages (Ashton 1964, 2014; Ashton and Hall 1992). Liana abundance results here from the outcome of recent decades of structural stability combined with, in places, the legacy of major destabilizing natural events in the past (Newbery et al. 1992, 1999). When such a forest is in the late stages of recovery after a historic major disturbance, a decline in liana abundance is expected (Campbell and Newbery 1993). The reasoning behind this postulate is that, as tree basal area abundance tends towards a site maximum, the main canopy closes, gaps become fewer and light conditions internal to the stand will be less favorable for liana recruitment and growth (Whitmore 1984).

With dry season seasonality and associated strong drought not being the issue, two factors remain though that could counteract an end-succession decline in SE Asian forests, namely a continuing rise in atmospheric carbon dioxide (Malhi and Phillips 2005), and perturbations caused by the intermittent effects of the El Nino Southern Oscillation (ENSO; Walsh 1996b, Wright 2005b) which open the canopy and lead the understorey to temporarily have more lighted and drier conditions (Walsh and Newbery 1999, Newbery et al. 2011). Lianas are also extensively distributed in logged forests of SE Asia (e.g. Pinard and Putz 1994), and have a major effect on forest regeneration; but that is a separate process from the dynamics of the mature primary forests, except for when these forests were last heavily disturbed by similar-scale natural events e.g. by extensive drought and/or fire (Beaman et al. 1985).

Wright et al. (2015) reported on changes in liana abundance at Pasoh, Peninsular Malaysia, between 2002 and 2014, in terms of canopy cover infestation on trees ≥ 30 cm dbh (95.4 cm gbh). They confirmed the lower susceptibility of dipterocarps to lianas compared with other tree families shown by Campbell and Newbery (1993). Tree mortality for all trees was affected when liana cover exceeded 75%. The change in proportions of all survivor trees infested was very small, declining from 52.3 to 47.9%. The frequency of trees with ≥ 50% cover of lianas barely altered. The samples were not independent though, because of the surviving trees in common.

An analysis of the change in liana infestation over a 30-year period in a late-successional primary lowland dipterocarp forest in N.E. Borneo is presented here to test expectations that liana infestation decreased over time but lianas were still having negative effects on tree growth and survival. The hypothesis of Campbell and Newbery (1993), that liana loss is largely controlled by branch fall for overstorey trees and whole tree death for understorey ones, is extended to show that this mechanism allows the dipterocarps, especially, to escape their parasitic liana load when reaching the upper canopy and becoming emergent.

## METHODS

### Study site

This study was conducted in two permanent plots of primary lowland dipterocarp rainforest within the Danum Valley Conservation Area (DVCA) in Sabah, north-eastern Borneo, Malaysia (Marsh and Greer 1992). The plots are each 4 ha (100 m × 400 m), lying parallel approx. 280 m apart and about 800 m west of the Field Centre (4°57’48’’N, 117°48’10’’E; Newbery *et al*., 1992). The area is ∼ 230 m above sea level, and both plots feature a ∼ 35-m change across gently undulating terrain. The climate is mostly aseasonal, with an average monthly temperature of 26.9°C and an average annual rainfall of 2832 mm (1986– 2007, Newbery *et al*. 2011). More detailed information is given in Newbery *et al*. (1992, 1996) and Walsh and Newbery (1999). The two plots are designated ‘main’ plots, MP1 and MP2, at Danum to distinguish them from 10 smaller neighbouring satellite ones.

The permanent plots were set up between 1985 and 1986 and have since been under ongoing long-term observation (Newbery *et al*. 1992). All trees with a girth at breast height ≥ 10 cm (gbh, at 1.3 m or above buttresses) were tagged, mapped, identified and gbh was measured. Since then, four more censuses have been made, the latest in 2015 (D. M. Newbery et al., unpubl. data). In each of those censuses, survivors, recruits and dead trees were recorded and gbh measured for all live trees.

The two plots had been initially intended as replicate areas of the forest locally, especially in terms of their species richness, general structure and biomass (Newbery et al. 1992). However, over time species’ tree size distributions diverged and the plots differed in their dynamics (Newbery et al. 1999, Lingenfelder and Newbery 2009; and unpubl. data). Thus, for the present liana study spanning 30 years, unless otherwise indicated, the plots were treated separately. Subsequently, this proved valuable as it led to much better insights into the liana dynamics than would have been achieved by a combined analysis which used them as simply statistical replicates.

### Fieldwork

From September 1987 to February 1988 and June to September 1989 (mid-date September 1988), a first liana census in the two plots was conducted by Campbell and Newbery (1993). Liana stems on all trees ≥ 30 cm gbh were counted (Zahnd 2018). The present study in June and July 2018, 30 years later, closely followed the procedure of the first one to achieve the highest feasible level of compatibility. Liana stems ≥ 2 cm gbh on all trees ≥ 30 cm gbh were again counted. Lianas were defined here as woody climbing plants which start growing from the ground. Lianas were regarded as individuals when they rooted close to the tree observed, or when crossing over into the target tree from another tree. Only lianas which had foliage in the canopy of the tree were counted: lianas which did not reach the crown of the tree, or only made a loop and descended, or that were clearly dead, were not counted. As in the 1988 census, lianas were not identified taxonomically because taking botanical samples for identification would have influenced the growth and mortality of the trees themselves undergoing long-term observation (Campbell and Newbery 1993).

For the present study tree girths measured in the first (1986) and last (2015) main plot tree censuses were applied to the liana counts from 1988 and 2018. These censuses gave the tree data closest in terms of plot census dates to those of the liana censuses. For both liana censuses, trees which died between 1986 and 1988 or between 2015 and 2018 respectively were disregarded, as were trees which only grew bigger than 30 cm gbh in the respective periods. By happenstance, the liana censuses were both at intervals of 2-3 years after the tree censuses, so if the dynamics in these relatively short periods can be taken as being equivalent there should have been very little bias introduced from the census matching. In this paper, data collected in 1986 and 1988 will be collectively referred to as ‘census 1’ (or ‘date 1’ in the analysis), and those collected in 2015 and 2018 as ‘census 2’ (‘date 2’).

The species codes in the 2017-digitized appendices of Campbell (1990), which formed the basis to the second liana census, were made fully compatible with the taxonomy of the last (fifth) 2015 plot census. Since the first census in 1986 (Newbery et al. 1992), some tree identifications to survivors were corrected (the more important ones mentioned in Newbery et al. 1999), with the consequence that for a few of them family memberships changed, and liana counts per tree in the different families reported in this paper may differ very slightly from those given by Campbell and Newbery (1993). Other tree names changed as a result of taxonomic revisions. Twenty-eight trees in four taxa changed names, but all remained in the same families. Within-plot elevations (ground height above plot origin) were available for all trees ≥ 30 cm gbh in 1986, and the 35 respectively 46 trees in MP1 and MP2 recruiting into this size class by 2015 were found by interpolation from plot coordinates (Lingenfelder and Newbery 2009). In the present liana study, the tree size class ‘small’ (10 -< 30 cm gbh) was not considered: trees ≥ 30 cm gbh are referred to as being medium sized, through large to very large.

## DATA ANALYSIS

In analyzing the changes in liana densities per tree between censuses 1 and 2, plots MP1 and MP2 were handled separately. Since the plots differed in topographic variation, tree size structure, forest dynamics and species composition, even in moderately small ways, combining them into one sample would have confounded several important and interesting differences that bear on the relationships between liana density and dynamics and these factors. Moreover, the plots are replicates at the 4-ha scale and separated enough spatially to provide some indication of forest variability on the local landscape. Apart from revisions to nomenclature, the 1988 liana data used here are the same as those reported in Campbell and Newbery (1993).

Trees at each census were divided into four size classes: 1, 30 -< 60; 2, 60 -< 120; 3, 120 -< 240; and 4, ≥ 240 cm gbh. These are referred to as medium-sized, large, very large and largest trees respectively: the class ‘small’ would be those 10 -< 30 cm gbh, not considered for the liana censuses. Over-understory index (OUI, scale 0 – 100), for the 100 most abundant species in the plots, was adopted from Newbery et al. (2011). It is based on first axis of a principal components analyses of ratios of numbers of trees ≥ 30 cm gbh/≥ 10 cm gbh and ratios of basal areas of the same size classes, the rescaled scores averaged over four plot tree censuses 1986 to 2007. Three storeys were designated: overstorey (OUI > 55), intermediate (OUI 20 – < 55) and understorey (OUI < 20). The 30-cm-gbh threshold is the same as the lower one for liana recording.

Analyses of the liana census data were mainly done in the computing language R (R_Core_Team. 2020; and Fox and Weisberg 2011). Special commands are listed in Appendix S5, where part 1 has the basic code for randomization testing of differences over time, and part 2 lays out the essential regression models, particularly those that allowed for spatial autocorrelation.

### Error distributions

The negative binomial distribution fitted the frequencies of number of lianas per tree far better than the Poisson, with χ^2^-values of 19.34 and 14.41 in 1988 for MP1 and MP2 respectively (df = 7), and correspondingly 13.89 and 14.50 in 2018 (df = 5). All four cases were nevertheless significant with *P* between 0.01 and 0.05. For comparison, fitting the Poisson χ^2^-values were 3344 and 1185 in 1988 and 2018 (plots together). The k-values (expressing degree of aggregation when < 1) were, for the four cases in order: 0.625, 0.491, 1.265 and 1.034; indicating that lianas were much more aggregated in 1988 than in 2018. The geometric Poisson (Polya-Aeppli) distribution gave very similar fits to the negative binomial, and since the latter is fairly widely incorporated as a family error distribution in most regression software, it was taken the error model. The remaining differences in fit were largely due to observed frequencies being less than expected ones for counts of 1 and 2 and the converse for counts of 3 and 4, for which no single-mode function can readily cater.

### Numbers of lianas

Negative binomial GLM regressions were made using the ‘glm.nb’ function in MASS library of R. Comparing fits (by anova) of nlianas ∼ ln(gbh) and nlianas ∼ ln(gbh) + elev for 1988 led to LR’s (log-likelihood ratios) of 11.86 (*P* ≤ 0.001) and 9.32 (*P* ≤ 0.01) in MP1 and MP2 respectively, and correspondingly for 2018 LR’s of 13.34 and 40.43 (both *P* ≤ 0.001), indicating that including elevation enhanced the model significantly. Here and later in model formulations, the term ‘nlianas’ is number of lianas per tree. Adding a ln(gbh)·elev interaction to the two-term model led to no improvement, LR’s of 0.14 and 0.49 in 1988 and 0.02 and < 0.01 in 2018 (all *P* > 0.10). By contrast, including instead a [ln(gbh)]^2^ term led in 1988 to LR’s of 3.10 (*P* ≤ 0.1) and 31.87 (*P* ≤ 0.001) in MP1 and MP2, and 2018 LR’s of 31.34 and 56.64 (both *P* ≤ 0.001). The final model was therefore: nlianas ∼ ln(gbh) + [ln(gbh)]^2^ + elev. Fitting the best model resulted in dispersion values of 0.948 and 1.001 in MP1 and MP2 for 1988, and 1.011 and 0.938 correspondingly for 2018. Diagnostic plots of Pearson residuals versus fitted showed reasonable spreads of points, with the typical striations for discrete data: residuals themselves however were positively skewed.

To test and cater for spatial autocorrelation h-likelihood GLM (HGLM) was run using the ‘hglm’ function in same-named hglm library of R, by including a SAR correlation-term based on plot inverse-distance matrices. As hglm does not cater for a negative binomial error, this had to be substituted by a quasi-poisson one; and models without the spatial autocorrelation adjustments compared with the negative binomial one. Running the selected model with the interaction also resulted in later being insignificant for all census x plot combinations. The method is described in Alam et al. (2015), Lee et al. (2017a, b) and Rönnegard et al. (2010).

### Tree survival

Tree survival between 1988 and 2018 was modelled with a logistic GLM regression (Hilbe 2009, 2011), as a standard ‘glm’ (binomial error) starting with survival ∼ sqrt(nlianas_88_). The transformation normalized the original nlianas variable to a large extent but not entirely. Adding ln(gbh_88_) and elevation resulted in varying degrees of improvement to the fits depending on the plot. With ln(gbh) the LR’s were 4.50 (*P* ≤ 0.05) and 2.67 (*P* > 0.10) in MP1 and MP2 respectively, whilst adding elevation led correspondingly to LR’s of 0.02 (*P* > 0.05) and 7.47 (*P* ≤ 0.01), although including both terms they were 4.58 (*P* ≤ 0.05) and 10.37 (*P* ≤ 0.01). As a compromise, in order to compare plots on a same-model basis, all three terms were retained to have a final model: survival ∼ sqrt(nlianas_88_) + ln(gbh_88_) + elev.

To account for spatial autocorrelation, an autologistic regression was run with the command ‘logistic.regression’ in the library spatialEco in R, using an inverse squared-distance matrix as a measure of correlation. Further, as an alternative to handling spatial autocorrelation affecting the model, repeated stratified sampling was achieved by taking one tree at random per 20-m × 20-m sub-plot (100 trees per plot), *N*’ = 500 times, and averaging the coefficients and statistics. This meant that trees on average 20-m apart were assumed to have independent liana counts. The repeated runs will have reused trees many times and not be independent, especially when there were very few per sub-plot from which to sample. This additional approach was intended to be confirmatory of the regressions, free of distance-decay function. Methods are outlined and discussed in Besag (1972), Cliff & Ord (1981), Augustin et al. (1996), Dormann (2007) and Zuur et al. (2009).

Analysis at the family level, for the Dipterocarpaceae and Euphorbiaceae, failed to converge to solution with the autologistic function in three of four cases, and it was replaced by a GLMM using the coordinates of 20-m × 20-m subplots as the cluster term in command ‘glmmML’ of the library glmmML in R (Rhodes et al. 2009, Brostrom and Holmberg 2011).

### Tree growth

With standard linear, or equivalently generalized least squares (GLS), regression (gaussian error) as the function ‘lm’ in R, relative growth rates (rgr, mm/m/yr) of trees, that is survivors, between 1988 and 2018, were fitted first as ∼ sqrt(nlianas_88_). Adding the term ln(gbh_88_) considerably improved model fits, with change in *F*-ratio (equivalent to LR) of 39.81 and 30.51 (both *P* ≤ 0.001) in MP1 and MP2 respectively, but not consistently so much in the case of elevation, the corresponding LR’s being 3.60 (*P* ≤ 0.1) and 0.62 (*P* > 0.10): both terms included led to combined improvements over sqrt(nlianas_88_) alone, LR’s of 21.08 and 15.53 (both *P* ≤ 0.001). Adding the squared size-term, [ln(gbh_88_)]^2^, to ln(gbh_88_) and sqrt(nlianas_88_) was of little gain, with *F*-ratios of 0.10 and 2.10, and adding to these elevation, 1.20 and 1.28 (all four at *P* > 0.10). Attempting to cater for some remaining non-normality in rgr, by using gamma or inverse-gaussian errors, resulted in scarce improvement in either fits or residual plots. Accordingly, the best parsimonious model was: rgr ∼ sqrt(nlianas_88_) + ln(gbh_88_).

With a rgr regression using the gaussian error, there were more ways of dealing with spatial autocorrelation than those (currently as software) available for regressions with binomial and negative binomial errors. Firstly, spatial dependence was modelled using a variogram of exponential form in GLS (a correlation structure), within the function ‘gls’ in library lnme in R. Comparing the accepted model form, without and with the spatial correlation term, led to modest improvement in the fits, with LR’s of 8.84 (*P* ≤ 0.05) and 5.54 (*P* ≤ 0.10) for MP1 and MP2 respectively. Using a spherical form in the variogram made hardly any difference. Randomization runs were conducted in the same way as they were for the survival data. A different, additional, approach was to use a correlation matrix again based on inverse distances but guided by prior analysis of spatial autocorrelation via a correlogram. Two indices, Moran’s I and Geary’s C, were calculated at 5-m interval distances between 0 and 40 m. Moran’s I for MP1 was significant at 5-10 m (*P* ≤ 0.001) and at 30-35 m (*P* ≤ 0.1), and for MP2 at 0-5 m (*P* ≤ 0.1), 10-15 and 15-20 m (both *P* ≤ 0.05). Geary’s C was significant in MP1 at 0-5 m (*P* ≤ 0.01) and 5-10 m too (*P* ≤ 0.05), and in MP2 again at 0-5 m (*P* ≤ 0.001) and 15-20 m (*P* ≤ 0.10). Thus, up to 20 m, both indices show significant spatial correlation, permitting use of the ‘spautolm’ function in library spatialreg in R, with SAR mode and range 0-20 m. The methods used here follow Pinheiro and Bates (2000), Haining (2010), Plant (2012), Bivand et al. (2013) and Borcard et al. (2018).

### Change in liana abundance with time

Changes in the proportion of trees carrying lianas, the frequencies of trees in liana density classes, and the mean number of lianas per tree, between 1988 and 2018 were estimated. Some of the trees were different ones at the two dates (those which died after 1988 were lost and those which recruited after 1988 to survive to 2018 were gained), but close to half were the same individuals (the survivors). (The 1721 survivors formed 51.2% of the trees in 1988 and 48.7% of those in 2018.) Therefore, the tree populations at the two dates were not fully statistically independent. Subsamples of different trees were accordingly selected for 1988 and for 2018, by taking at random half of those in 1988 (a uniform distribution of the tag numbers), flagging them, and the unflagged trees contributed to the sample for 2018. Some unflagged trees will not have survived 1988 to 2018, and they were replaced (approximately in terms of numbers) by recruits appearing in 2018. Thus no one tree was present in both dates’ subsamples. Statistical tests were repeated *N*’ = 500 times as a Monte Carlo procedure (Manly 1997). This randomization procedure not only eliminated temporal dependence but it also reduced the effects of spatial autocorrelation.

Differences in the proportions of trees having lianas, that is presence or absence of lianas, at the two dates were tested using Pearson’s chi-squared statistic (2 × 2 test; df = 1). For frequencies of trees in liana density classes (0, 1, 2…r lianas) the chi-squared statistic was again applied (2 × r test, df = r − 1), with the provision, achieved by pooling higher tail classes, that no class had < 5 trees. Firstly, all trees were used in single tests, without any subsampling. Randomization tests were then run *N*’ times, the mean statistic found and its significance, and the null hypothesis was rejected overall when ≥ 80 % of the randomization runs were individually significant at the *P* ≤ 0.05. This procedure admits a power of 0.8 and type II error rate of β = 0.2 (Cohen 1988, 1992). The mean chi-squared values were taken as representing ‘typical’ differences in proportions free of any temporal autocorrelation effects. Additionally, McNemar’s chi-squared test of symmetry (Agresti 2007), was employed to ask whether proportions of trees with or without lianas changed with date, for just the 1988-survivors. The symmetry statistic could only be calculated for counts in the density classes when taking all trees: the two main families had insufficient numbers.

Change in proportions of trees with lianas between the two dates could be analysed too with the negative binomial GLM, to ask whether mean numbers per tree differed significantly, and whether the relationship between proportion and size of tree changed over time. Regressions were run with subsamples at the two dates from the same *N*’ = 500 randomizations; they compared nested models 1 to 4: nlianas ∼ year, ∼ year + ln(gbh), ∼ year * ln(gbh), and ∼ year * ln(gbh) + [ln(gbh)]^2^ + year: [ln(gbh)]^2^. The last two models tested whether differences in proportions were further accountable for by linear and non-linear interactions between year and ln(gbh). Interpreting the interactions in GLMs between a categorical factor (e.g. year) and continuous variables (ln[gbh] and [ln(gbh)]^2^), and then using the term to infer estimates of the expected response variable (nlianas) requires special consideration. Guidance to the calculations are found in Hilbe (2009), Cameron and Trivedi (2013) and Long and Freese (2014).

### Liana abundance and dynamic status

Starting with the trees censused in 1988, proportions of trees with lianas could be compared between those dying and those surviving by 2018 with the chi-squared test of association, and the numbers of lianas per tree similarly compared with negative binomial GLM. In complementary way, those trees at 2018 that were survivors from 1988 and those that had recruited (into the ≥ 30 cm gbh population) by then could also be compared.

### Liana densities per tree for individual species

Mean number of lianas per tree were found for the 51 and 52 species in 1988 and 2018 respectively with ≥ 20 trees per species, from fitting individual negative binomial distributions. Over- understorey index (OUI) values were already available for the 100 most abundant species. Six further species occurred in a 100-list for 2015 and their OUI-values were taken as the means of the genera to which they belonged. Mean gbh per species was derived from the main plot census records. Stem rgr in girth of small trees (10 -< 50 cm gbh) for period one, between the first two tree censuses (1986-1996), have been reported in Newbery et al. (1999), though the 100 most abundant species (N./plot ≥ 10 cm gbh) available for here are the very slightly revised values of 2010, and were for only ‘valid’, i.e. unproblematic, gbh estimates (Lingenfelder and Newbery 2009). In this way 45 and 47 (of the 51 and 52) species could be matched with rgr-values averaged across the two plots.

## RESULTS

### Background tree dynamics

At census 1 (1986|1988) there were 1704 and 1659 trees of ≥ 30 cm gbh in MP1 and MP2 respectively (together 3363), and by census 2 (2015|2018) correspondingly 1699 and 1832 (3531), discounting the few that died between matched tree and liana censuses. Between the censuses, 926 and 716 (together 1642) trees died in MP1 and MP2, and correspondingly 921 and 889 (1810) trees recruited into the size class. The full data set consisted then of 3363 + 1810 = 5373 trees involved in both censuses. The annualized mortality rates were accordingly 2.58 and 1.87%, and the annualized recruitment rates 1.45 and 1.33%, in MP1 and MP2 respectively. (Mortality was based on the proportions of the numbers of trees alive in 1986 dying by 2015, whilst recruitment was based on proportions of trees alive in 2015 which had joined the population since 1986; see Newbery and Lingenfelder 2009 for Danum). Thus, mortality rate was higher than recruitment in both plots, and both mortality and recruitment rates in MP1 were higher than their corresponding values in MP2, especially for mortality (38% higher). In the 30 years MP1 had a 26% higher turnover than did MP2 (2.02 versus 1.60%). Overall, trees ≥ 30 cm gbh occurring in both censuses and both plots belonged to 355 species, 143 genera and 55 families. The frequency distributions of tree gbh were very similar between plots and dates, especially close for MP1 (Appendix S1: Fig. S1).

### Liana frequency distributions

The nine to ten most abundant tree families (Table 1 and Appendix S1: Table S1) combined made up 75% and 71% of all trees in MP1 and MP2 respectively. The most abundant families were the *Dipterocarpaceae* and the *Euphorbiaceae*, followed by the *Meliaceae*, *Lauraceae*, *Phyllanthaceae* and *Annonaceae*. Relative abundances of the first 13 families per plot plus all other pooled into a 14^th^, did not change significantly between dates (MP1: χ^2^ = 9.57, df = 13, *P* = 0.73; MP2: χ^2^ = 18.05, df = 13, *P* = 0.16), nor did the overall 10-cm gbh classed distributions of all trees (pooling trees ≥ 150 cm) between years (MP1: χ^2^ = 14.19, df = 12, *P* = 0.286; MP2: χ^2^ = 9.35, df = 12, *P* = 0.67; see also Appendix S1: Fig. S1). This largely rules out that changes in overall liana density were due to major shifts in the tree community over the 30 years. Nevertheless, MP1 and MP2 did differ in one important respect: MP1 had more very large trees than MP2 (Appendix S1: Fig. S1), although they decreased in number more in MP2 than MP1 between 1988 and 2018. These very large trees were mostly dipterocarps.

**Table 1.** Number of trees, relative abundance, proportion of liana infested trees and average number of lianas per tree for all trees and those in the two most abundant families, in each census in main plots MP1 and MP2.

| Plot | Tree family | Number of trees<br>per plot |  | Trees with lianas<br>(%) |  | Average number of<br>lianas per tree |  |
| --- | --- | --- | --- | --- | --- | --- | --- |
|  |  | 1988 | 2018 | 1988 | 2018 | 1988 | 2018 |
| MP1 | All trees | 1704 | 1699 | 60.3 | 65.0 | 2.164 | 1.686 |
|  | Dipterocarps | 246 | 247 | 45.9 | 40.1 | 1.841 | 1.000 |
|  | Euphorbs | 206 | 190 | 64.6 | 68.9 | 1.922 | 1.889 |
| MP2 | All trees | 1659 | 1832 | 54.3 | 56.7 | 2.017 | 1.333 |
|  | Dipterocarps | 330 | 353 | 36.7 | 36.8 | 1.361 | 0.841 |
|  | Euphorbs | 219 | 225 | 53.0 | 53.3 | 1.995 | 1.053 |

In total, 2865 and 2442 liana stems were counted in plots 1 and 2, respectively. This is a 21.8% and 27.0% decrease since the first census (3688 and 3347 lianas in plots 1 and 2, respectively). The mean number of lianas per tree, excluding trees without any lianas, decreased in MP1 and MP2 from respectively 2.16 and 2.02 in the first census to 1.69 and 1.33 in the second one, decreases of 21.8 and 34.2% (Table 1). For all trees, the proportion of liana-infested trees significantly increased by 4.7% in MP1 and by 2.4% in MP2. None of the major families individually showed significant changes in the proportions, although in MP1 trees in the Dipterocarpaceae decreased whilst those in the Euphorbiaceae correspondingly increased: differences for these two families in MP2 were negligible (Table 1).

Frequency distributions of counts of lianas per tree declined smoothly in plots MP1 and MP2, with relatively more trees with no or few (1 to 3) lianas, and relatively fewer trees with many (≥ 7) lianas in 2018 than in 1988 (Fig. 1a, b). The distributions appeared approximately exponential in form. For the Dipterocarpaceae inflation of zero counts was obvious, with the difference between years matching the overall one in MP2 but not so well in MP1 (Fig. 1c, d). The Euphorbiaceae followed a scaled down representation of the overall plot distributions in MP2 (also for the zero-class), but MP1 not (Fig. 1e, f). The two plots appeared on first examination to be showing different dynamics in liana densities over time, which justified them being separately analyzed statistically. The liana distributions for 1988 are the same as those used in Campbell and Newbery (1993: Figs 2 and 3), albeit presented here slightly differently. In that earlier paper the size distributions of trees in the Dipterocarpaceae and Euphorbiaceae were compared (ibid.: Fig. 1).

**Fig. 1.**
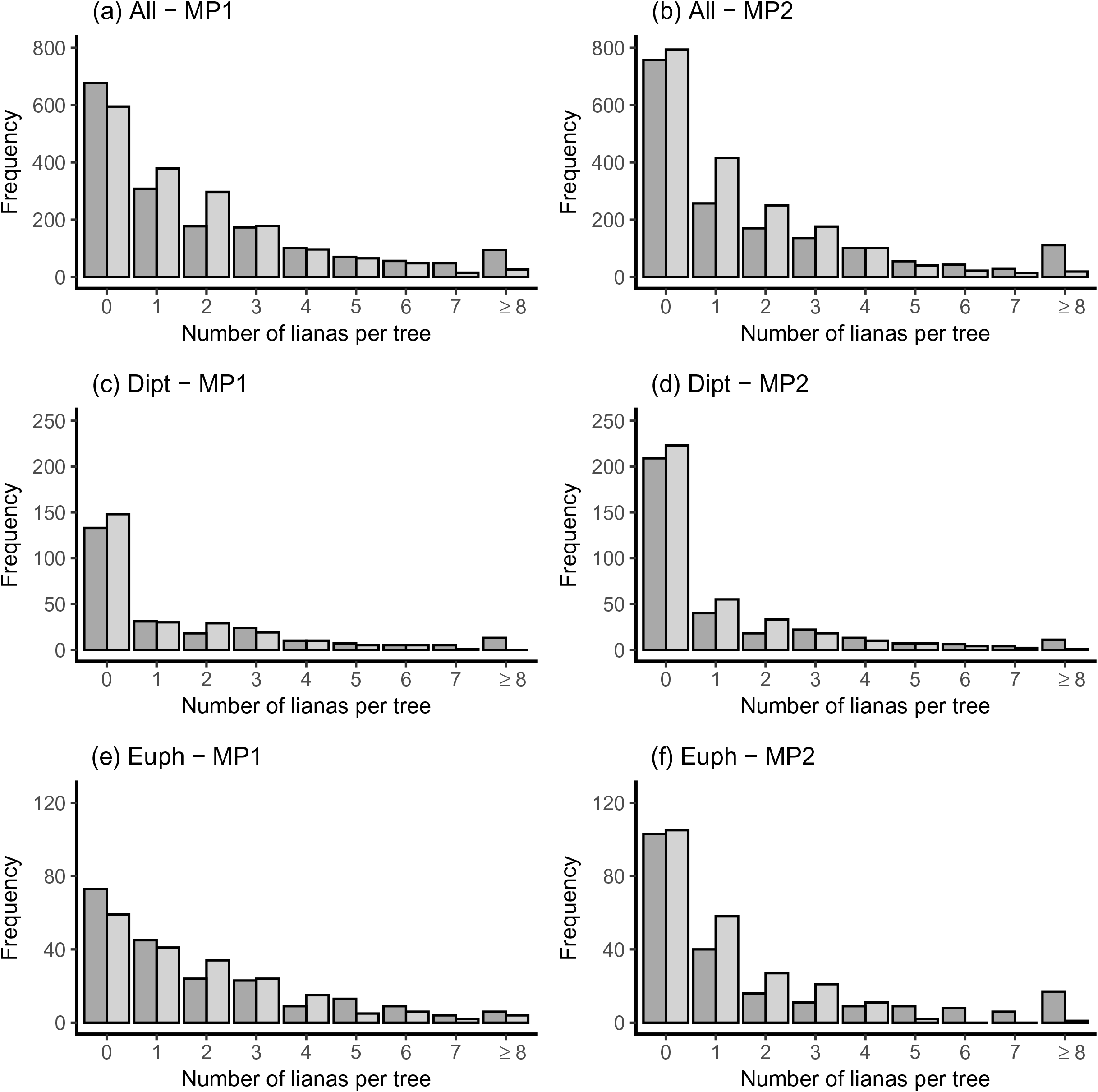
Frequency distributions of trees in classes of numbers of lianas per tree in 1988 (dark grey bars) and in 2018 (light grey bars) in the two main plots MP1 and MP2 at Danum, for trees in (a, b) all families, (c, d) Dipterocarpaceae, and (e, f) Euphorbiaceae.

**Fig. 2.**
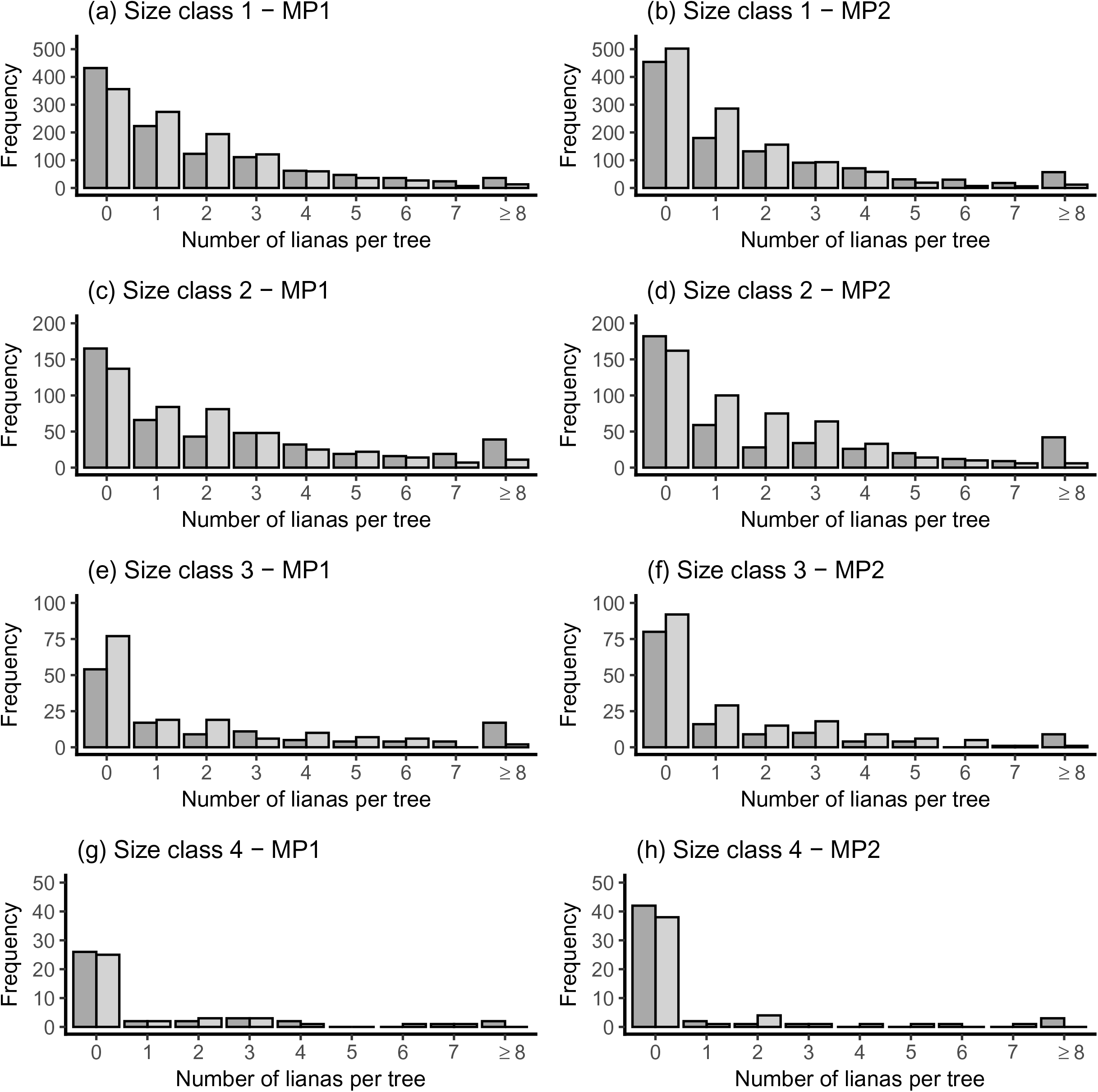
Frequency distributions of trees in classes of numbers of lianas per tree in 1988 (dark grey bars) and in 2018 (light grey bars) in the two main plots MP1 and MP2 at Danum, for trees in four tree girth size classes (gbh): (a, b) 1, 30 - < 60 cm; (c, d) 2, 60 - < 120 cm; (e, f) 3, 120 -< 240 cm; and 4, ≥ 240 cm.

**Fig. 3.**
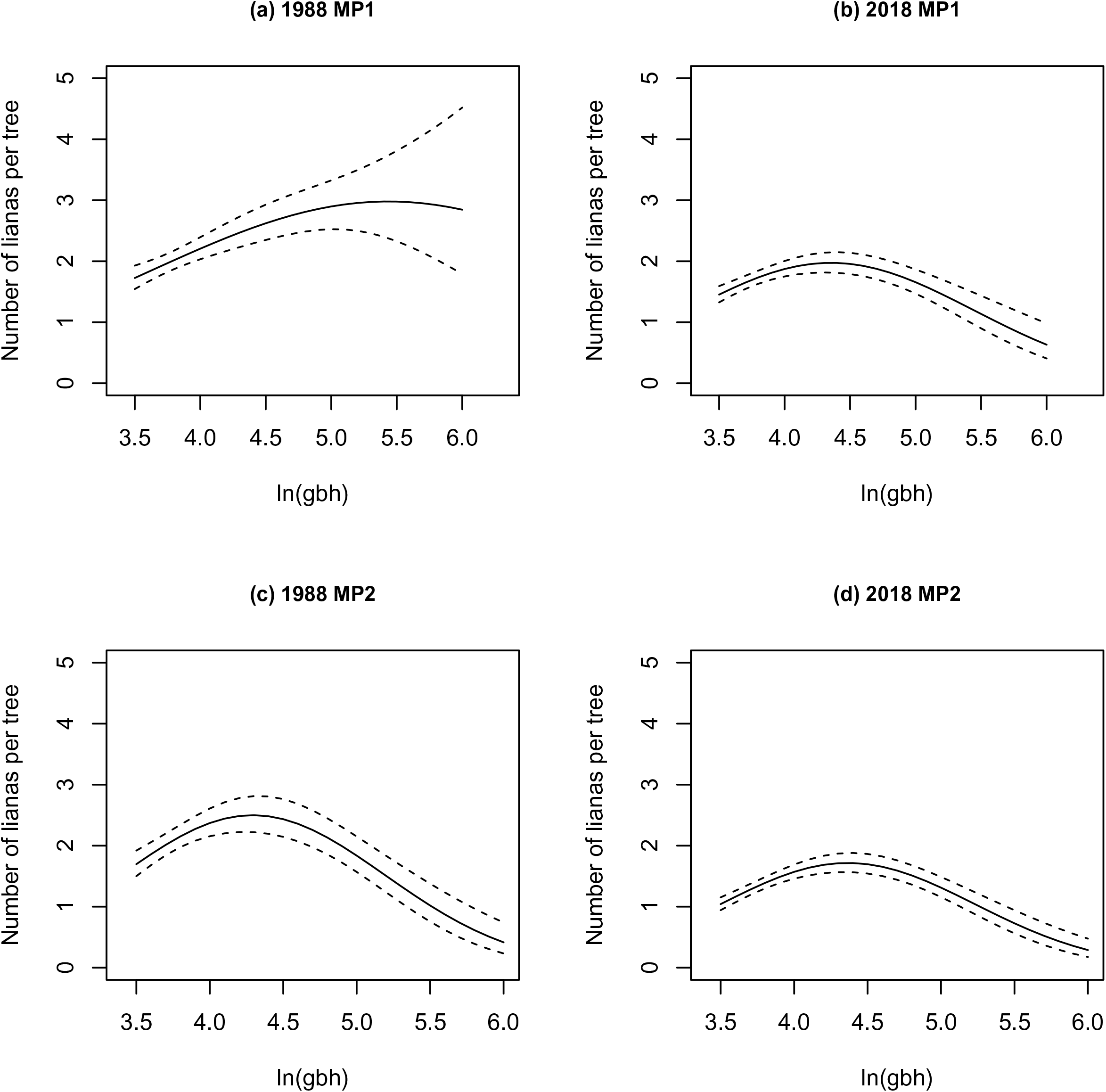
Fitted relationships from negative binomial GLM regressions for the number of lianas per tree versus tree tree girth at breast height size (as ln[gbh]) in main plots 1 and 2, at each census date: solid line, 1988; dashed line, 2018.

Considering the frequency distributions of numbers of lianas per tree in the four size classes (Fig. 2), the gradual decline for 0 to ≥ 8 lianas per tree, in both plots and at both dates, in the medium-sized trees (class 1) becomes increasingly more discontinuous as proportionally more trees up to the very large trees (class 3) have no lianas, and among the largest (class 4), zero counts become dominant. This implies indirectly that as the trees become larger they are shedding their lianas.

### Number of lianas and tree size

The relationship between number of lianas per tree and tree size (gbh) could be described by a quadratic form, for MP1 in 1988 rather weakly, but more strongly in 2018 (Table 2, Fig. 3). For MP2, the fits were similarly strongly quadratic at both census dates. More lianas were recorded on medium than on small and large trees: the distinction between censuses in MP1 appeared mainly due to large trees in 1988 having similar loads to medium-sized ones. Consequently, change in numbers over time was more marked for MP1 than for MP2. In 1988, coefficients were statistically weaker for MP1 than for MP2; in 2018 they were highly significant for both plots. Adjustment for spatial autocorrelation was small although its effect was more noticeable for MP1 in 1988, especially on the gbh-terms. Regression using HGLM with the quasi-Poisson error gave similar results to the negative binomial GLM (Appendix S2: Tables S1a, b), suggesting that the variance-mean relationship in numbers of lianas was close to linear (see ver Hoef and Boveng 2007).

**Table 2.** Regressions of numbers of lianas per tree on tree girth at breast height (gbh) and elevation (HGLM, quasi-Poisson error) in the censuses of 1988 and 2018, with a term, rho, for spatial autocorrelation (SAR^†^).

| census | term | MP1<sup>‡</sup> |  | MP2 |  |
| --- | --- | --- | --- | --- | --- |
| | | est ± se | t | est ± se | t |
| 1988 | intercept | $-2.493 \pm 1.451$ | $-1.719^{\circ}$ | $-9.938 \pm 1.843$ | $-5.391^{***}$ |
| | ln(gbh) | $1.365 \pm 0.668$ | $2.043^{*}$ | $5.097 \pm 0.867$ | $5.877^{***}$ |
| | ln(gbh)<sup>2</sup> | $-0.1284 \pm 0.0762$ | $-1.684^{\circ}$ | $-0.5987 \pm 0.1007$ | $-5.944^{***}$ |
| | elevation | $-0.01512 \pm 0.00676$ | $-2.238^{*}$ | $-0.01218 \pm 0.00399$ | $-3.051^{**}$ |
| 2018 | intercept | $-6.825 \pm 1.480$ | $-4.612^{***}$ | $-11.620 \pm 1.615$ | $-7.195^{***}$ |
| | ln(gbh) | $3.540 \pm 0.695$ | $5.097^{***}$ | $5.669 \pm 0.761$ | $7.453^{***}$ |
| | ln(gbh)<sup>2</sup> | $-0.4092 \pm 0.0808$ | $-5.064^{***}$ | $-0.6477 \pm 0.0885$ | $-7.316^{***}$ |
| | elevation | $-0.01041 \pm 0.00349$ | $-2.981^{**}$ | $-0.01541 \pm 0.00247$ | $-6.248^{***}$ |

<sup>†</sup> $\rho$ : MP1<sub>88</sub>, 0.0943; MP1<sub>18</sub>, 0.0381; MP2<sub>88</sub>, 0.1224; MP2<sub>18</sub>, 0.0604.
<sup>‡</sup> in 2018 two observations with high leverage omitted.
\*\*\*, $P \leq 0.001$ ; \*\*, $P \leq 0.01$ ; \*, $P \leq 0.05$ ; $^{\circ}$ , $P \leq 0.10$ , <sup>ns</sup>, $P > 0.10$ .

Pseudo-*R*^2^-values (as %), based on differences in deviances for fitted and null models (in the latter the mean was the sole term), and their random and fixed components (in parenthesis), were in 1988, 24.2 (22.0, 2.2) and 16.8 (14.8, 2.0) for MP1 and MP2 respectively, and correspondingly in 2018, 20 (17.4 and 2.6) and 10.2 (4.6, 5.6). The statistically weaker terms for MP1 in 1988 match with the highest, whereas the strongest terms in MP2 in 2018 had the lowest, random component. Even so, the fixed component was small at just 2-6%. The model fits are descriptive, and not fully hypothesis testing, because ln(gbh) and [ln(gbh)]^2^ were not statistically independent variables (i.e. some collinearity was present). It follows that the absolute values of the *t*-statistic associated with the ln(gbh) and [ln(gbh)]^2^ coefficients will be similar. The high random component (meaning a large between-tree variance) in the MP1-1988 model might have been responsible for the less pronounced quadratic fit (Appendix S1; Fig. S2).

Of the eight combinations of plot x census x family two showed significance for the ln(gbh) and [ln(gbh)]^2^ terms: Dipterocarpaceae in MP2 for 1988 and Euphorbiaceae in MP1 for 2018 (Appendix S2: Table S1c). Accounting for spatial autocorrelation (HGLM) affected the coefficients only very slightly but the significances more, being lower for the Euphorbiaceae (remaining at *P* ≤ 0.05). Elevation was otherwise also significant for two other combinations (*P* ≤ 0.05). Trees at the family levels were less dense than overall, so the influence of spatial autocorrelation was less important.

### Tree survival

Tree survival was significantly negatively related to number of lianas in both plots, although weakly with gbh and inconsistently (positively) with elevation in MP2 (Table 3). Including a spatial correlation term (in the autologistic regression) changed the slope of survival on number of lianas very little (Appendix S2: Table S2a), even though the spatial covariate was significant for MP1 (Table 3). The coefficient for gbh was likewise minimally affected. Stratified random sampling gave average slopes slightly smaller than those for the autologistic fit, especially for the number of lianas term in MP1 (Appendix S2: Table S2b). Since number of lianas in the regression were those recorded in 1988, it is reasonable to conclude that increasing liana loads reduced tree survival in the 30 years. Despite the clear trend, the pseudo-*R*^2^ values were very low, 0.8 and 1.5% in MP1 and MP2 respectively. At the family level, survival was significantly negatively dependent on number of lianas for euphorbs in both plots (*P* ≤ 0.05) and dipterocarps in MP2 (*P* ≤ 0.01), although for MP1 the slope was in the same direction (Appendix S2: Tables S2c, d). Interestingly, the slopes of all four relationships are higher than those for all species in the plots (Table 3). What the liana loads were later after 1988 on trees that died by 2018 is unknown.

**Table 3.** Auto-logistic regressions of tree survival 1988-2018 on number of lianas per tree (nlianas) and tree girth at breast height (gbh) in 1988, and elevation (GLM, binomial error), accounting for spatial autocorrelation (covariate).

| term | MP1 |  | MP2 |  |
| --- | --- | --- | --- | --- |
| | est ± se | *t* | est ± se | *t* |
| intercept | $-0.834 \pm 0.380$ | $-2.20^*$ | $-0.434 \pm 0.385$ | $-1.13^{\text{ns}}$ |
| nlianas<sup>1/2</sup> | $-0.1454 \pm 0.0482$ | $-3.02^{**}$ | $-0.1947 \pm 0.0482$ | $-4.40^{***}$ |
| ln(gbh) | $0.2035 \pm 0.0914$ | $2.23^*$ | $0.1583 \pm 0.0901$ | $1.76^{\text{ns}}$ |
| elevation | $-0.0024 \pm 0.0061$ | $-0.39^{\text{ns}}$ | $-0.0120 \pm 0.0044$ | $-2.74^{**}$ |
| covariate | $0.0638 \pm 0.0309$ | $2.06^*$ | $0.0678 \pm 0.0454$ | $1.49^{\text{ns}}$ |

\*\*\*, $P \leq 0.001$ ; \*\*, $P \leq 0.01$ ; \*, $P \leq 0.05$ ; °, $P \leq 0.10$ , <sup>ns</sup>, $P > 0.10$ .

### Tree growth

The negative effects of number of lianas on tree relative growth rate (rgr_88-18_) were highly significant (*P* ≤ 0.001). Elevation was non-significant, but gbh more significant (also negative) than number of lianas (Table 4). Coefficients were very similar for MP1 and MP2. Accounting for spatial autocorrelation led to scarcely any change in coefficient values, for either the two regression approaches or for stratified random sampling, indicating that local spatial dependencies were very weak (Appendix S2: Tables S3a-c). Nevertheless, testing for autocorrelation avoided type I errors. The pseudo-*R*^2^-values for growth models were much higher than for survival, 22.5 and 32.0% for MP1 and MP2.

**Table 4.** General least squares regressions of tree relative growth rate (rgr) 1988-2018 on number of lianas per tree (nlianas) and tree girth at breast height (gbh) in 1988 (GLS, mixed-model, gaussian error), accounting for spatial autocorrelation (corExp variogram).

| term | MP1 |  | MP2 |  |
| --- | --- | --- | --- | --- |
| | est ± se | *t* | est ± se | *t* |
| intercept | 0.747 ± 0.071 | 10.544*** | 0.603 ± 0.060 | 10.047*** |
| nlianas<sup>1/2</sup> | -0.0369 ± 0.0095 | -3.878*** | -0.0353 ± 0.0083 | -4.261*** |
| ln(gbh) | -0.0998 ± 0.0170 | -5.875*** | -0.0755 ± 0.0143 | -5.288*** |

\*\*\*, $P \leq 0.001$ ; \*\*, $P \leq 0.01$ ; \*, $P \leq 0.05$ ; °, $P \leq 0.10$ , <sup>ns</sup>, $P > 0.10$ .

Simple negative binomial regressions of number of lianas in 1988 on rgr had slopes of −0.574 ± 0.201 (*z* = −2.853, *P* ≤ 0.01) and −0.742 ± 0.220 (*z* = −3.376, *P* ≤ 0.001) for MP1 and MP2. Repeating for number of lianas now in 2018 on rgr the corresponding slopes were −0.577 ± 0.159 (*z* = −3.643, *P* ≤ 0.001) and −0.508 ± 0.172 (*z* = −2.958, *P* ≤ 0.01). Regressing the number of lianas in 1988, with a fuller model that included ln(gbh) and [ln(gbh)]^2^ of 1988, and elevation, the coefficient for rgr decreased for MP1 and increased moderately for MP2 (Appendix S2: Table S4). A similar regression with gbh of 2018 instead, led to the rgr coefficients increasing much more for both plots, so that by partialling out the effects of gbh in the negative dependence on liana load on rgr increased. Lastly, the *change* in number of lianas between 1988 and 2018, treated as a Poisson variable (*n* + 22 — to make all changes positive) was not significantly related to rgr in MP1 (−0.006 ± 0.028, *z* = 0.234, *P* = 0.82) and in MP2 (0.024 ± 0.027, *z* = 0.879, *P* = 0.38). Adding ln(gbh) and elevation terms did not improve the fitting overall or significance of rgr.

Linear regression of rgr on number of lianas and ln(gbh) at the family level showed the expected dependence on size for the dipterocarps but not the euphorbs (Appendix S2: Table S3d, e) but largely no or weak significance of dependence on number of lianas. The use of the variogram-based spatial autocorrelation adjustment scarcely affected the results: with coefficients increasing slightly, and for euphorbs in MP2 a small rise in significance (still only *P* ≤ 0.05).

### Changes in liana abundance – contingency analysis

Using all trees at both census dates, there was a small significant (*P* < 0.01) increase of almost 5% in the proportion of trees with lianas between 1988 and 2018 for MP1 but not MP2 where it was less at ca. 2.5% (Table 5). At the family level there were no clear significant differences, largely because sample sizes were much smaller than for all plot trees. The randomization testing supported the result for MP1 but only at *P* < 0.05, and 80 percentile values were all insignificant (Table 5). Here a qualification is needed: χ^2^ values were positively skewed, more so in MP2 than MP1, so that the medians were lower than the means. This made the significance for all trees in MP1 possibly more marginal. Considering just the survivors, the change in MP1 and also in MP2 was larger with 8-9% increases over time, and more strongly significant (*P* < 0.001) than when deaths after 1988 and recruits at 2018 were involved. Between families, there was pronounced difference in MP1: with a strong decrease in proportion of trees in the Dipterocarpaceae (not quite significant however) counterbalanced by a much larger increase for the Euphorbiaceae (∼22%). These differences roughly mirror the smaller relative changes at the ‘all trees’ level. However, for MP2 dipterocarps did not alter proportions and euphorbs increased just slightly (Table 6). Therefore, over all trees the proportion with lianas increased slightly in the 30 years, more so for just survivors, but mostly for surviving euphorbs in MP1.

**Table 5.** Changes in the proportion of liana infested trees between censuses using Pearson’s χ^2^ test on all the trees and based on the randomized subsampling (N’ = 500). The mean and 80^th^ percentile of χ^2^-values is given for the randomizations. McNemar’s tests were calculated using only the survivors. All tests had 1 degree of freedom.

| plot | family | all trees |  | randomization |  | survivors<br>(McNemar) |  |
| --- | --- | --- | --- | --- | --- | --- | --- |
| | | $\Delta\text{prop. (\%)} \pm$<br>95 CI | $\chi^2$ | Mean $\chi^2$ | 80% $\chi^2$ | $\Delta\text{prop.}$<br>(%) | $\chi^2$ |
| MP1 | All trees | $+4.7 \pm 3.306$ | 7.861** | 4.210*<sup>†</sup> | 1.611<sup>ns.</sup> | +8.9 | 15.890*** |
| | Dipterocarps | $-5.8 \pm 9.131$ | 1.493<sup>ns.</sup> | 1.136<sup>ns.</sup> | 0.076<sup>ns.</sup> | -8.5 | 2.161<sup>ns.</sup> |
| | Euphorbs | $+4.3 \pm 9.777$ | 0.669<sup>ns.</sup> | 0.674<sup>ns.</sup> | 0.012<sup>ns.</sup> | +22.2 | 6.050* |
| MP2 | All trees | $+2.4 \pm 3.358$ | 1.852<sup>ns.</sup> | 1.484<sup>ns.</sup><sup>‡</sup> | 0.189<sup>ns.</sup> | +7.5 | 13.499*** |
| | Dipterocarps | $+0.1 \pm 7.396$ | <0.001<sup>ns.</sup> | 0.410<sup>ns.</sup> | 0.008<sup>ns.</sup> | 0.0 | 0.000<sup>ns.</sup> |
| | Euphorbs | $+0.3 \pm 9.649$ | <0.001<sup>ns.</sup> | 0.425<sup>ns.</sup> | 0.001<sup>ns.</sup> | +2.9 | 0.098<sup>ns.</sup> |

<sup>†</sup>, <sup>‡</sup>: medians (ranges), MP1 3.70 (0.0 – 18.6), MP2 0.95 (0.0 – 10.4)
\*\*\*, $P \leq 0.001$ ; \*\*, $P \leq 0.01$ ; \*, $P \leq 0.05$ ; °, $P \leq 0.10$ , <sup>ns.</sup>, $P > 0.10$ .

**Table 6.** Changes in the frequency of trees in liana density classes between censuses using Pearson’s χ^2^ test on all the trees and based on the randomized subsampling (N’ = 500). The mean and 80^th^ percentile of χ^2^ values is given for the randomizations. McNemar’s tests were calculated using only the survivors. Sample sizes for individual family McNemar’s tests were too small.

| plot | family | all trees |  | randomization |  |  | survivors<br>(McNemar) |  |
| --- | --- | --- | --- | --- | --- | --- | --- | --- |
| | | df | $\chi^2$ | df | mean $\chi^2$ | 80% $\chi^2$ | df | $\chi^2$ |
| MP1 | All trees | 9 | 101.981*** | 8 | 53.015*** | 45.245*** | 15 | 43.9*** |
|  | Dipterocarps | 6 | 14.270* | 4 | 7.016<sup>ns.</sup> | 3.879<sup>ns.</sup> |  |  |
|  | Euphorbs | 7 | 9.441<sup>ns.</sup> | 4 | 4.159<sup>ns.</sup> | 2.062<sup>ns.</sup> |  |  |
| MP2 | All trees | 10 | 134.211*** | 8 | 68.690*** | 59.357*** | 15 | 67.515*** |
|  | Dipterocarps | 6 | 14.267* | 4 | 7.668<sup>ns.</sup> | 4.518<sup>ns.</sup> |  |  |
|  | Euphorbs | 5 | 41.228*** | 4 | 16.304** | 11.907* |  |  |

\*\*\*, $P \leq 0.001$ ; \*\*, $P \leq 0.01$ ; \*, $P \leq 0.05$ ; °, $P \leq 0.10$ , <sup>ns.</sup>, $P > 0.10$ .

Considering counts of lianas per tree is more informative. Taking all trees (i.e. all families), and the single χ^2^-tests with no randomized subsampling, very highly significant differences (*P* < 0.001) in the relative frequency distributions between dates are highlighted for both plots (Table 6, Fig. 1). The large χ^2-^values in general were close to normally distributed, and so the medians were also almost the same as the means. Not only do the randomization tests support the simpler ones, but also so do the McNemar’s symmetry tests for just survivors. Figure 1 shows clearly the change in class frequencies with a decline in numbers of trees with very high loads (≥ 8 lianas per tree), and an increase in trees with 1-3 lianas per tree. Considering the families separately, the randomization tests did not hold up the confidence in the single overall χ^2^-test (the latter significant at *P* < 0.05 in both plots, Table 6) for the dipterocarps. Conversely, whilst the Euphorbiaceae showed no evidence of differing class proportions in MP1, they did in MP2, supported by the randomization tests (*P* < 0.01).

The failure to detect significance in the dipterocarps under the randomization may have been because there were few trees in very-large tree size class, the one with the high liana loads especially in 1988 and hence sample variances increased when taking only half of the trees at each census. Fig. 1 (c, d) does not show large differences within classes, especially the case for MP1. However, coming to the Euphorbiaceae, (Fig. 1 e, f) there is a marked difference between dates within classes, and especially for MP2 where the frequency classes 6, 7 and ≥ 8 are empty in 2018 (cf. the numbers in 1988), and the increases in classes 1-3 similar to the overall plot level frequencies. Evidently, the changes in liana abundance between 1988 and 2018, while minimal in terms of proportions of trees with any lianas and none, are happening in the redistribution of the frequency classes: fewer to no trees with very many lianas and more trees with a few lianas by 2018. That trend was happening more strongly in MP2 than MP1, and on account largely of changes within the Euphorbiaceae.

### Changes in liana abundance – regression analysis

Negative binomial GLM regressions using the randomized subsamples involved four models, and for all trees each of them is shown in Table 7a. Model 3, with three terms (aside from the intercept), is interesting because, for MP1, year, size and their interaction were all significant. Without the interaction term, year became more significant but size less so, though more noticeable was that between models 2 and 3 the sign of the coefficient for year changed from negative to positive and accordingly trends with ln(gbh) were catered for more in the latter by the interaction term. For MP2 it was just year that had a consistent and strong (negative) effect. Model 3 was expanded to model 4 to have two further terms, [ln(gbh)]^2^ and its interaction with year, to accommodate the quadratic element of the relationship nlianas and ln(gbh) and thus the different shapes of the curves between years particularly for MP1. Even with more terms in model 4, than the other models, individual terms were all less significant for MP1 but for MP2 ln(gbh) and [ln(gbh)]^2^ featured strongly (Table 7a). It is important to recall that model 4 was more a descriptive than hypothesis-testing model.

**Table 7.** Negative binomial regression of number of lianas per tree on census date (year, as a factor) and tree girth at breast height (as ln[gbh]) for (a) all trees (all four models), and (b) those in the families Dipterocarpaceae and Euphorbiaceae (just model 1). The four models are: 1, nlianas ∼ year; 2, nlianas ∼ year + ln[gbh]; 3, nlianas ∼ year + ln[gbh] + year:ln[gbh]; and 4, nlianas ∼ year + ln[gbh] + year:ln[gbh] + (ln[gbh])^2^ + year:(ln[gbh])^2^. The values are averages of N’ = 500 randomizations.

| (a) |  |  | MP1 |  | MP2 |  |
| --- | --- | --- | --- | --- | --- | --- |
| M |  | terms | est ± se | z | est ± se | z |
| 1 | All | intercept | 0.771 ± 0.044 | 17.422*** | 0.701± 0.049 | 14.258*** |
|  |  | year<sub>18</sub> | −0.249 ± 0.064 | −3.901*** | −0.412± 0.070 | −5.903*** |
| 2 | All | intercept | 0.040 ± 0.237 | 0.169<sup>ns</sup> | 0.596 ± 0.254 | 2.348* |
|  |  | year<sub>18</sub> | −0.240 ± 0.064 | −3.774*** | −0.413 ± 0.070 | −5.913*** |
|  |  | ln[gbh] | 0.180 ± 0.057 | 3.112** | 0.026 ± 0.062 | 0.425<sup>ns</sup> |
| 3 | All | intercept | −0.492 ± 0.327 | −1.502<sup>ns</sup> | 0.831 ± 0.356 | 2.337* |
|  |  | year<sub>18</sub> | 0.930 ± 0.471 | 1.974* | −0.874 ± 0.503 | −1.735• |
|  |  | ln[gbh] | 0.310 ± 0.080 | 3.874*** | −0.032 ± 0.087 | −0.368<sup>ns</sup> |
|  |  | year<sub>18</sub>:ln[gbh] | −0.290 ± 0.116 | −2.508* | 0.114 ± 0.123 | 0.924<sup>ns</sup> |
| 4 | All | intercept | −3.778 ± 2.095 | −1.781<sup>o</sup> | −10.756 ± 2.543 | −4.209 *** |
|  |  | year<sub>18</sub> | −3.636 ± 3.110 | −1.168<sup>ns</sup> | −1.681 ± 3.661 | −0.464<sup>ns</sup> |
|  |  | ln[gbh] | 1.862 ± 0.973 | 1.888<sup>o</sup> | 5.449 ± 1.192 | 4.549 *** |
|  |  | year<sub>18</sub>:ln[gbh] | 1.860 ± 1.451 | 1.281<sup>ns</sup> | 0.506 ± 1.718 | 0.300<sup>ns</sup> |
|  |  | (ln[gbh])<sup>2</sup> | −0.180 ± 0.111 | −1.584<sup>ns</sup> | −0.636 ± 0.138 | −4.597 *** |
|  |  | year<sub>18</sub>:(ln[gbh])<sup>2</sup> | −0.248 ± 0.167 | −1.488<sup>ns</sup> | −0.047 ± 0.199 | −0.244<sup>ns</sup> |
| (b) |  |  | MP1 |  | MP2 |  |
| M |  | terms | est ± se | z | est ± se | z |
| 1 | Dipt | intercept | 0.599 ± 0.161 | 3.747*** | 0.302 ± 0.155 | 1.959• |
|  |  | year | −0.607 ± 0.234 | −2.589** | −0.481 ± 0.221 | −2.172* |
| 1 | Euph | intercept | 0.648 ± 0.116 | 5.590*** | 0.687 ± 0.137 | 5.035*** |
|  |  | year | −0.019 ± 0.168 | −0.116<sup>ns.</sup> | −0.647 ± 0.202 | −3.207** |

\*\*\*, $P \leq 0.001$ ; \*\*, $P \leq 0.01$ ; \*, $P \leq 0.05$ ; °, $P \leq 0.10$ , <sup>ns</sup>, $P > 0.10$ .

Model 4 was overall the best-fitting model for MP1 and MP2 but for slightly differing reasons. Averaged over the 500 iterations, of all four models, AIC was lowest in model 4 for both plots, and model 4 reached highest pseudo-*R*^2^ values in MP1 and MP2 (2.64 and 4.56 % respectively) (Appendix S3: Table S1). Highest LRs were attained when model 4 was compared to model 1, again in both plots (LR_03_ = 32.3 and 46.4); the changes from model 1 to 3 were nearly as significant (LR_13_ = 22.7 and 45.8). The six paired comparisons were the models coded 12, 13, 14, 23, 24 and 34. Notable was that LRs for models 1 to 2 in both plots were relatively low (Appendix S3: Table S1), indicating that model fitting markedly improved when interactions between year and ln(gbh) and [ln(gbh)]^2^ were estimated. Probabilities of the χ^2^ change in deviance when comparing models were lowest for model 1 versus 4 in MP1, and even lower for models 1, 2 and 3 with model 4 in MP2.

Considering the frequencies with which models were selected across the iterations, lowest AIC-values were attained with model 4 in 497 of 500 iterations for MP1 (model 3 in three), and all 500 for MP2. Likewise, model 4 had the highest LR-values among all six comparisons in all 500 for both plots, and similarly pseudo-*R*^2^ was highest for all 500 per plot. Such consistency is not surprising given the small SEs of the statistics (Appendix S3: Table S1). Delta-AIC was found for models paired in the same way done for LR, and here 475 of 500 were largest for model 1 minus model 4 for MP1, but 373 of 500 for model 3 minus model 4 in MP2 (Appendix S3: Table S2). Similarly, the significance of the χ^2^ measure of deviance change was highest for 455 or 500 iterations in MP1 for the 14 comparison and 438 or 500 in MP2 for the 34 one. LR was never highest for the model 23 comparison or were the corresponding probabilities of the deviance change the lowest; very rarely were these statistics the highest respectively lowest for model comparisons 12 and 13 either (Appendix S3: Table S2). These last results point to an important difference between the plots: whilst model 4 was overall best for both, the most improvement for MP1 was between model 1 and 4, but for MP2 it was between models 3 and 4.

In these glm.nb fits, year was coded as a factor. This meant that year_18_ was the difference from the reference year_88_, and likewise interactions between ln(gbh) and [ln(gbh)]^2^ with year were differences in their corresponding slopes (Appendix S2: Table S4). For MP1 nlianas in 2018 became a decreasing proportion of the nlianas in 1988 as gbh increased, going from 85% at ln(gbh) = 3.5 (gbh = 33.1 cm) to 40% at ln(gbh) = 5.5 (gbh = 244.7 cm) (Appendix S2: Table S5a). But for MP2 the change was very different, from 62 rising to 73% for the same gbh range (Appendix S2: Table S5b). So whilst model 4 was the better one for MP1 (cf. models 1-3) and showed the strongest change with respect to gbh, its main terms became less, and the interaction terms conversely more, significant.

Model 4 was also run on the full data, i.e. all trees without random subsampling at the dates, and the coefficients fitted matched the averages from the randomizations very well. (Appendix S2: Table S6a). Further, the coefficients from model 4 could be used to reconstruct the individual equations for the dependence of nlianas on ln(gbh) and [ln(gbh)]^2^ at each date (similar to those reported in Table 2 without the elevation term, again matching closely (Appendix S2: Table S6b). This demonstrated that in testing for change in nlianas over time (the year factor) the gbh curves were being modelled by the interaction terms correctly. Predicted (marginal) means of nlianas in 1988, and in 2018 — using a mean ln(gbh) and [ln(gbh)]^2^ common to years — were very similar across the four models, with just small decreases of 1.5-1.8% for MP1, and 2.8-3.0% in MP2, between models 1 and 4 (Appendix S2: Table S7a). Predicted means at the family level were also very similar (Appendix S2: Table S7b). The means resulting from models 1 and 2 are, naturally, virtually the same as the values given in Table 1.

### Liana abundance and dynamic status

Among the trees that died 1988-2018, a higher proportion had lianas present in 1988 compared with those that survived (Table 8). This was especially significant in MP2. Correspondingly, dead trees had on average more lianas per tree than survivors. Trees in both the Dipterocarpaceae and Euphorbiaceae followed the same trends, again more strongly in MP2 than MP1. Survivors of dipterocarps had less than half the numbers per tree than those that died. Recruits on the other hand by 2018 had proportionally fewer trees with lianas than survivors, and less lianas per tree, especially in MP1 (Table 9). Differences were less strong at the family level, except for dipterocarps where the proportion of trees with lianas was significantly less on survivors than on recruits in both plots, though numbers per tree were similar.

**Table 8.** The proportions of trees with lianas for those trees that survived versus died (*n_s_*, *n_d_*) between census 1 and 2 (1988, 2018), and their corresponding mean numbers per tree, in the two plots for all families (all trees) and for those trees in the Dipterocarpaceae (Dipt) and Euphorbiaceae (Euph). χ^2^ is from the Pearson test of association, *z* is the statistic from the binomial GLM regression.

| Family | Plot | Number of trees | | % of trees with lianas | | $\chi^2$<sup>†</sup> | Mean number lianas per tree | | |
| --- | --- | --- | --- | --- | --- | --- | --- | --- | --- |
| | | die | survive | die | survive | | die | survive | $z$<sup>‡</sup> |
| All | MP1 | 926 | 778 | 62.7 | 57.3 | 4.96* | 2.326 | 1.972 | -2.367* |
|  | MP2 | 716 | 943 | 59.6 | 50.3 | 14.03*** | 2.402 | 1.725 | -4.238*** |
| Dipt | MP1 | 105 | 141 | 50.5 | 42.6 | 1.22<sup>ns</sup> | 2.114 | 1.638 | -1.059<sup>ns</sup> |
|  | MP2 | 133 | 197 | 44.4 | 31.5 | 5.14* | 2.000 | 0.929 | -3.215** |
| Euph | MP1 | 152 | 54 | 69.7 | 50.0 | 5.95* | 2.112 | 1.389 | -2.078* |
|  | MP2 | 114 | 105 | 61.4 | 43.8 | 6.10* | 2.509 | 1.438 | -2.538* |

<sup>†</sup>: df = 1; <sup>‡</sup> df = 1, $n_s+n_d-2$ .
\*\*\*, $P \leq 0.001$ ; \*\*, $P \leq 0.01$ ; \*, $P \leq 0.05$ ; <sup>ns</sup>, $P > 0.10$ .

**Table 9.** The proportions of trees with lianas for those that survived versus recruited (*n_s_*, *n_r_*) between census 1 and 2 (1988, 2018), and their corresponding mean numbers per tree, in the two plots for all families (all trees) and for those trees in the Dipterocarpaceae (Dipt) and Euphorbiaceae (Euph). In comparing Tables 8 and 9, the numbers of trees surviving are the same (as in Table 8), but the numbers of trees dying and recruiting differed (column ‘rec−die’ here). χ^2^ is from the Pearson test of association, *z* is the statistic from the binomial GLM regression.

| Family | Plot | Number of trees |  | % of trees with lianas |  |  | Mean number lianas per tree |  |  |
| --- | --- | --- | --- | --- | --- | --- | --- | --- | --- |
| | | recruit | rec–die | survive | recruit | $\chi^{2\dagger}$ | survive | recruit | $z^\ddagger$ |
| All | MP1 | 921 | –5 | 66.2 | 64.0 | 0.84<sup>ns</sup> | 1.923 | 1.486 | –4.536*** |
|  | MP2 | 889 | 173 | 57.8 | 55.5 | 0.93<sup>ns</sup> | 1.408 | 1.253 | –1.906<sup>o</sup> |
| Dipt | MP1 | 106 | 1 | 34.0 | 48.1 | 4.42* | 0.950 | 1.066 | 0.512<sup>ns</sup> |
|  | MP2 | 156 | 23 | 31.5 | 43.6 | 4.99* | 0.716 | 1.000 | 1.692<sup>o</sup> |
| Euph | MP1 | 136 | –16 | 72.2 | 67.6 | 0.19<sup>ns</sup> | 2.093 | 1.809 | –0.822<sup>ns</sup> |
|  | MP2 | 120 | 6 | 46.7 | 59.2 | 3.03<sup>o</sup> | 0.943 | 1.150 | 1.144<sup>ns</sup> |

$^\dagger$ : df = 1; $^\ddagger$ df = 1, $n_s+n_r-2$ .
\*\*\*, $P \leq 0.001$ ; \*, $P \leq 0.05$ ; <sup>o</sup>, $P \leq 0.10$ , <sup>ns</sup>, $P > 0.10$ .

### Liana densities per tree for individual species

The mean numbers of lianas per tree for the 51 and 52 more frequent tree species in MP1 and MP2 together at 1988 and 2018 were merged to 61 species in common (Appendix S4: Table S1). Mean species’ number of lianas per tree was very weakly correlated with OUI (*r* = −0.006, df = 49, *P* = 0.97) and ln(gbh) (*r* = −0.047, df = 49, *P* = 0.74) in 1988 though slightly more strongly in 2018 (*r* = −0.175, df = 50, *P* = 0.21; *r* = −0.278, df = 50, *P* = 0.046 correspondingly). OUI and ln(gbh) were highly correlated in 1988 (*r* = 0.859, df = 48, *P* < 0.001; with *Nothaphoebe* sp. dropped because its ln(gbh) was very heavily outlying) and in 2018 (*r* = 0.904, df = 50, *P* < 0.001). However, the poor correlations hide an interesting separation between six to seven canopy/emergent dipterocarps, with high OUI-values but very low numbers of lianas per tree, versus the rest that follow a general positive trend (Figs. 4 and 5). The separation is weak in 1988 but becomes distinct in 2018. Similar patterns arise when plotting against ln(gbh) instead of OUI. This separated group of dipterocarps clearly have very much lower numbers of lianas per tree than would be predicted from the general trend with increasing OUI of all the others species.

**Fig. 4.**
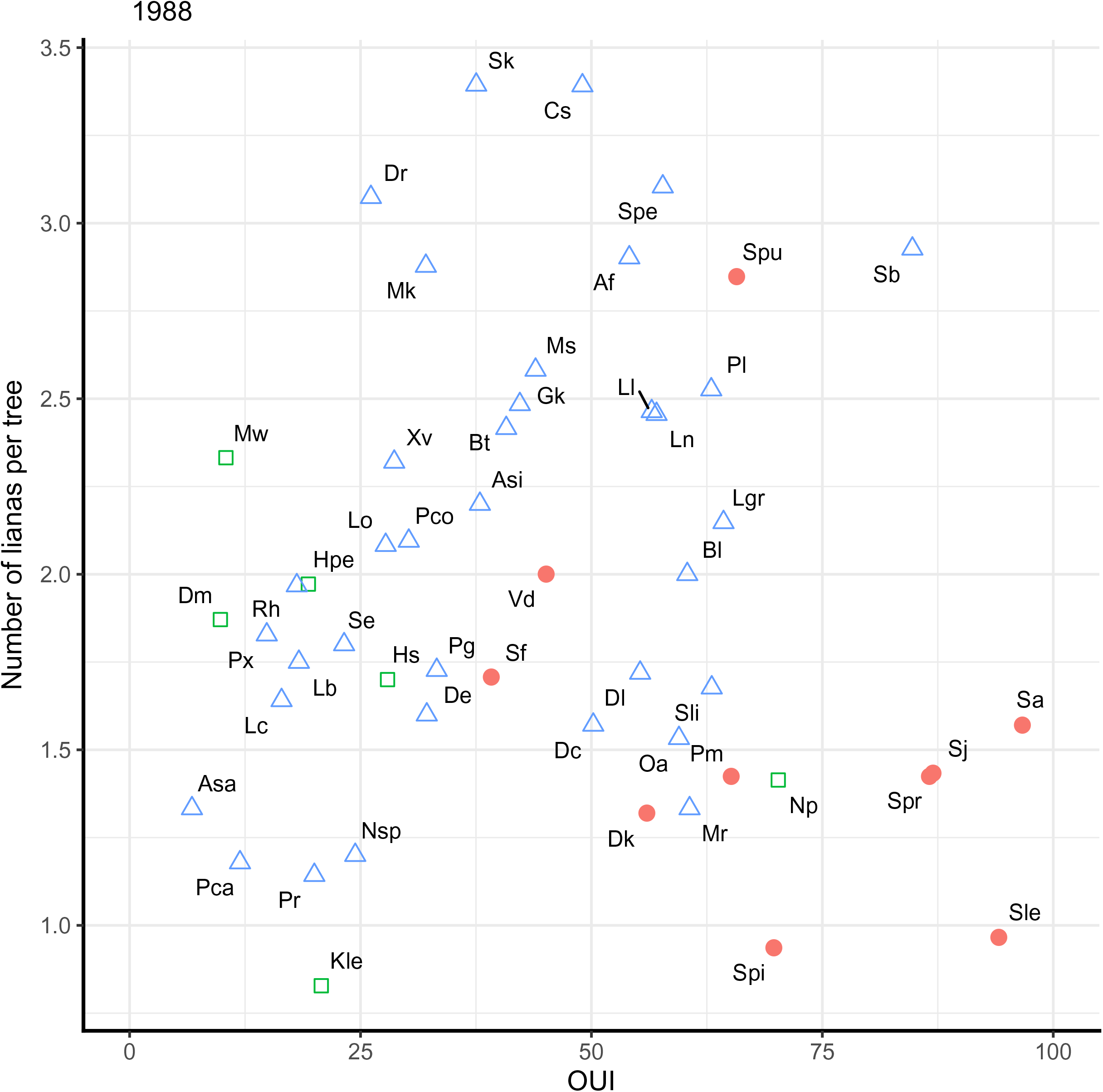
Relationships between mean species’ number of lianas per tree and the over-understorey index (OUI), for species with ≥ 20 trees in plots MP1 and MP2 combined in 1988. Closed red circles, Dipterocarpaceae; open blue squares, Euphorbiaceae; open green triangles, other families.

**Fig. 5.**
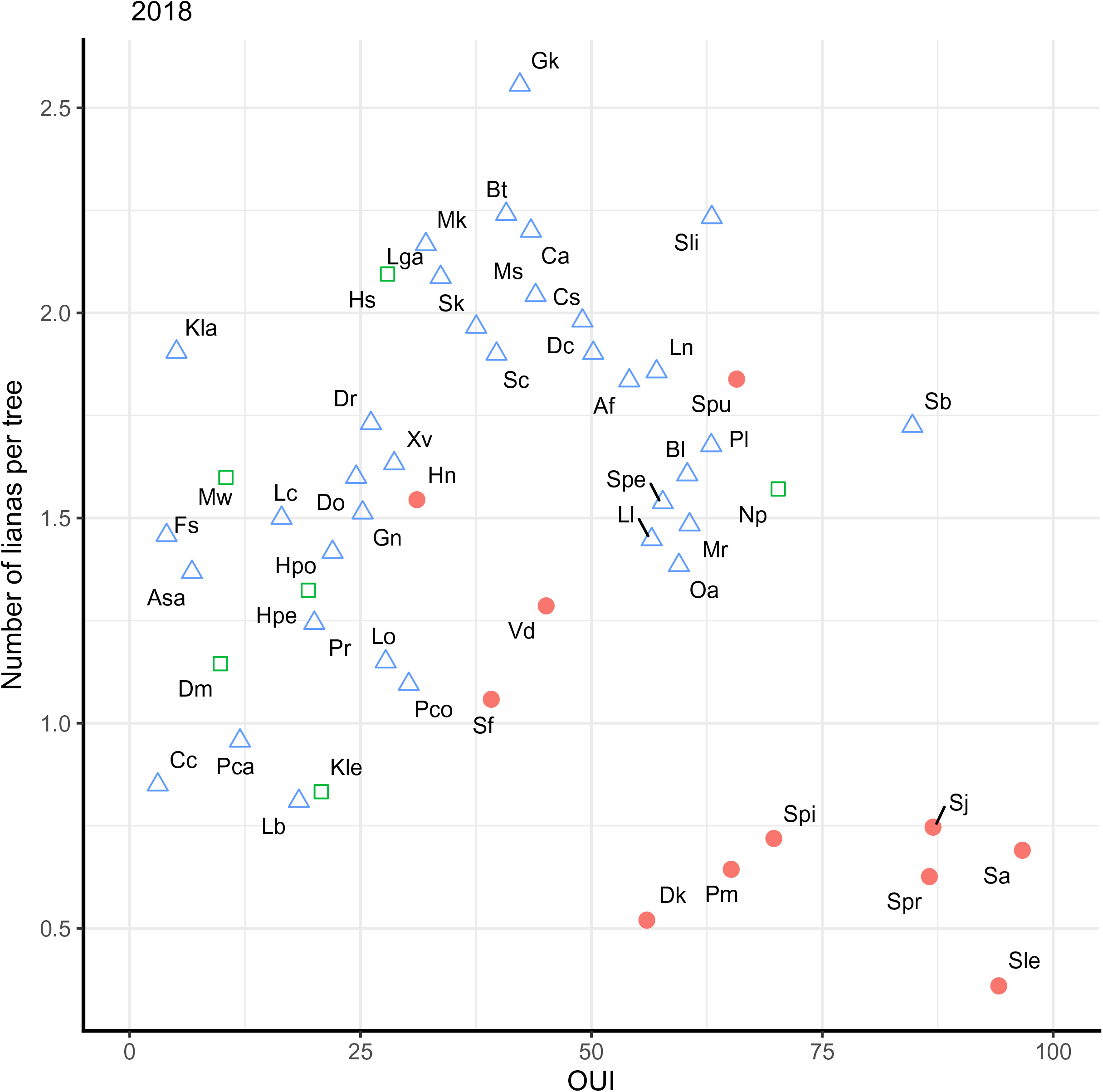
Relationships between mean species’ number of lianas per tree and the over-understorey index (OUI), for species with ≥ 20 trees in plots MP1 and MP2 combined in 2018. Symbols as in Fig. 4.

Species’ means of numbers of lianas per tree were weakly correlated with percentages of 1988 trees that died by 2018 (%*d*) at both 1988 (*r* = −0.103, df = 49, *P* = 0.48) and 2018 (*r* = −0.061, df = 50, *P* = 0.67), and also this was the case for correlations with ln(rgr) of small trees in 1988 (*r* = −0.094, df = 48, *P* = 0.52; no rgr-estimate for *Nothaphoebe sp*.) and 2018 (*r* = −0.269, df = 50, *P* = 0.054). Separation of groups, especially of the large dipterocarps was much less evidence although the 6-7 species that featured on the OUI graphs also had among the highest *rg*r-values when their trees are small (Appendix S4: Fig. S1).

Forty-two species occurred at both dates on the ≥ 20-trees criterion, and their liana numbers per tree in 1988 and 2018 were positively correlated (*r* = 0.661, df = 40, *P* < 0.001; Fig. 6). Although the variables at the two dates are not fully independent from one another (they shared survivors and their lianas), the dependence of numbers in 2018 on 1988 is instructive even if the significance level attached is inflated due to some autocorrelation. nlianas_18_ = 0.411 + 0.511·nlianas_88_ (*F* = 31.1, df = 1, 40; *P* < 0.001). On average, all species had fewer lianas in 2018 than 1988, but the reduction was most when the numbers in 1988 were higher. The fitted line shows the difference from a 1:1 expectation. The average of the plots in 1988 was 2.091 lianas per tree, and 1.510 per tree in 2018 (from Table 1). Putting this average for 1988 in the fitted linear model gave an estimated mean of 1.472. An interesting feature of Fig. 6 is that most of the dipterocarps that attain very large sizes are below the fitted line, having among the lowest means in 2018, reflecting the patterns in Fig. 5. The exception again is *S. pauciflora*. Differences between species’ means of numbers of lianas per tree (1988- minus 2018-values) were moreover also not significantly correlated with ln(rgr) of small trees (*r* = 0.093, df = 40, *P* = 0.56) or with %*d* (*r* = −0.115, df = 40, *P* = 0.47). No phylogenetic adjustment was necessary since the results were at best only marginally significant. Liana density was related to species mean size (ln[gbh]), and form of size distribution defining storey (OUI) when conditioned on tree family, i.e. dipterocarps vs non-dipterocarps, but not with the basic growth rate of their trees when small and species mortality rates.

**Fig. 6.**
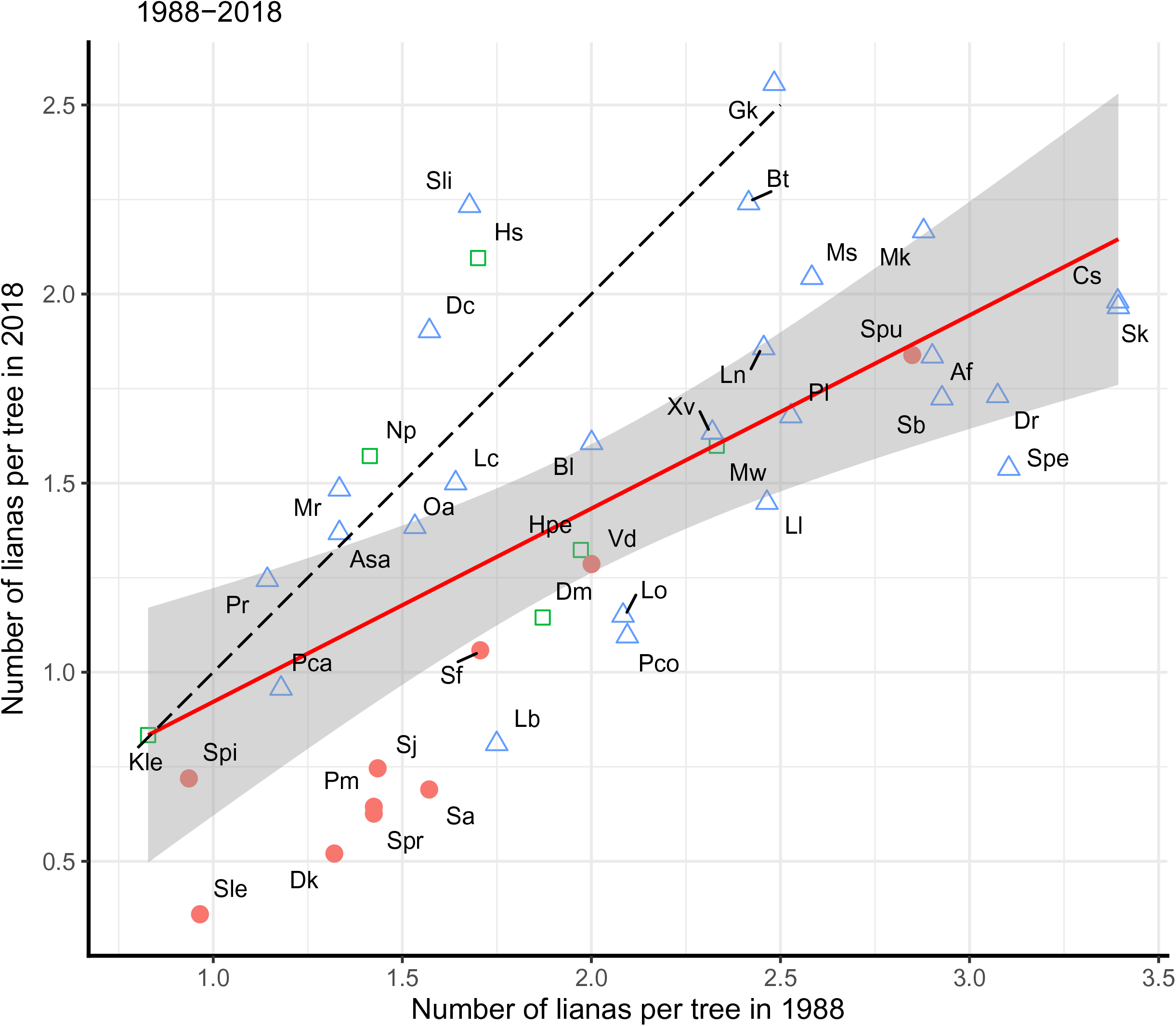
Differences in species’ mean numbers of lianas per tree (1988- minus 2018-values) for species with ≥ 20 trees in both censuses, in plots MP1 and MP2 combined. Symbols as in Fig. 4. The red line is the linear fit for all points, with 99% confidence band; the black dashed line is the 1:1 no-change expectation.

## DISCUSSION

### Changing liana abundance

The two liana censuses covered a recent 30-year period in a forest considered to be in a stage of late successional recovery from a major natural drought disturbance, probably in the 1870’s. Indicators were the very small proportion of plot area with gaps, very few pioneer-stage trees, few large lying stems, no evidence of major wind damage, and total tree basal area approaching an expected site maximum accompanied by declining tree density (Newbery et al. 1992). This contrasts strongly with the majority of liana studies to date which were conducted in heavily disturbed and secondary successional forests, or forests with a strong component of fast-growing light-demanding species (Visser et al. 2018a).

The closest and most relevant comparison to the present study is that of Wright et al. (2015). At Danum the proportion of trees ≥ 30 cm dbh (95.4 cm gbh) with lianas present was 44.5% in 1988 and 48.6% in 2018, values slightly less on average than at Pasoh (see Introduction), those at Pasoh decreasing and at Danum increasing a little. If the crown position class ‘emergent’ for Pasoh (Fig. 1c of Wright et al. 2015, for 2014) can be equated to the size-class 4 at Danum (Fig. 3, for 2018), the corresponding proportions of the very largest trees with no lianas were c. 25% and 75%, a three-fold higher liana-free stem frequency at Danum. Further comparisons are problematic because the studies used different methods of recording liana abundance (% canopy cover vs numbers per stem), and had different census intervals (12 vs 30 years). With some approximate adjustments, the rate of decline seemed much steeper at Danum than at Pasoh.

Between the two censuses tree size frequency distributions and major family contributions did not change significantly, although above a gbh of 240 cm tree frequencies at Danum were more erratic with numbers of trees in 2018 larger than in 1988 for the two largest classes (Appendix S1: Fig. S1). The average numbers of lianas per tree decreased by 25% in the 30-year period, though the proportions of trees with lianas increased by 3.5%. This came about because in 2018 relatively more trees had few lianas (1-3 in number), and relatively fewer trees had many lianas (≥ 8), than in 1988. Numbers of trees with no lianas remained on average very similar. The two plots differed importantly, however, in their liana dynamics. Large-sized trees had mainly higher liana loads than medium-sized and very large ones, except for MP1 in 1988 when very large trees had similarly high loads to large trees. The statistical errors were relatively high here due to the fewer trees in the very large-tree size class. Accordingly, this latter class showed a much faster decline between the censuses in MP1 than in MP2. Number of lianas per tree decreased significantly with topographic elevation overall: they were fewer on ridges than lower slopes. Despite the often high significance of coefficients from the model fitting, total variances accounted for were low, indicating that many other unknown sources of variation besides tree size and elevation were important.

In both plots trees with many lianas in 1988 survived significantly less well than those with none or few. Tree survival also decreased with elevation, more strongly in MP2 than MP1, but it was unrelated to tree size. Likewise, when liana loads increased this led to significantly reduced rgr of gbh, but rgr was unaffected by elevation. Final model fits accounted for far more variance in growth (27% on average) than in survival (∼ 1%). Thus, if lianas were affecting tree growth negatively, then survival would be expected to decrease as well, as reduced growth normally enhances tree mortality. Partialling out elevation still left survival and growth depending on tree size. Differences between families in the dependence of their trees’ growth rates on liana load were not consistent between plots though. For dipterocarps the relationship was negative in MP2 but not so in MP1, and for euphorbs it was slightly negative both in MP1 and in MP2. The decrease in rgr with increasing tree size for all trees was highly significant, for the dipterocarps too, but not for the euphorbs, at the family level.

Monte Carlo randomization testing showed that the increase in proportions of trees with lianas was marginally significant in MP1 but not significant in MP2. For survivors alone the changes were stronger and more significant, indicating perhaps that mortality and recruitment of trees in the period was offsetting accumulation on survivors. The most robust results, statistically, came from the randomization testing for numbers of lianas per tree, confirming the decline between 1988 and 2018 in MP1 and MP2, in the proportion of trees with high numbers of lianas per tree, and the corresponding increase in the proportion of trees with very few lianas. This form of testing, taking independent tree samples at the two census dates (i.e. samples having different survivors), also removes ‘survival bias’ due to trees with heavy liana loads often dying along with their loads and the liana counts becoming zero (Visser et al. 2018a, b). This bias has, *inter alia*, important consequences for understanding how liana loads, or ‘burdens’, affect tree population growth rates (Muller-Landau and Vissier 2019). Randomization also avoided the more essential problem of a lack of independence caused by survivors being counted in the samples of both census dates. As to be expected for a long-lived structural parasite, liana numbers increased with the age of the surviving host unless they were shed. Even so, trees die for many other reasons, and infestation by lianas is but one contributor.

The weak, or lacking, differences in mean numbers of lianas between censuses in the Dipterocarpaceae in both plots may indicate that for this family, summed across the size classes, lianas were being gained at a similar rate to them being lost without undue mortality of large and very large trees. As discussed later, the interaction between lianas and the very largest trees points to an important process operating in this dominant tree family. Most of the large to very large trees in the Danum plots are dipterocarps (Newbery et al. 1992, 1996). For the smaller Euphorbiaceae, however, there was little difference in mean number of lianas in MP1 yet substantial decreases in MP2. The negative binomial regressions for the changes in numbers of lianas per tree, under randomization testing, furthermore highlighted the importance of the year x size interaction, which again differed between plots. The hierarchical model comparisons confirmed that largest reductions over time were for the very large trees, especially in MP1. The differing patterns in small-scale dynamics of the trees and their lianas highlights the fundamental role of location, a spatial variability between plots set in the context of larger-scale temporal change.

In this study only the number of lianas per tree stem were recorded and not liana stem diameter or canopy crown infestation. At two other sites, these variables were found to be rather weakly correlated with one another (van der Heijden et al. 2008, Cox et al. 2019), and that might have been the case too at Danum. Good reliable observations of all parts of crowns of all trees (i.e. understorey, intermediate and overstorey) from the forest floor was not feasible at Danum because of the high tree density and the strongly gradated forest profile. The lack of good viewing points would have meant omitting an unacceptably high proportion of the trees sampled and a resulting selective bias in cover values. Number of lianas per tree is a first approximation to liana abundance: a more quantitative measure would have been liana basal area per tree. How lianas were distributed above in trees, and indeed which ones climbed from medium to large individuals was not recorded. Putz and Chai (1987) found though that each canopy liana (> 2 cm dbh) connected on average 1.4 trees > 20 cm dbh in primary dipterocarp forest at Lambir, Sarawak.

When considering rgr response, the causal direction is not unambiguous. On the one hand, high liana loads might have reduced rgr more than did low ones but, on the other, trees that were growing fast, may have discouraged the survival of lianas more than those growing slowly. This might be connected with branch shedding: faster growing trees would probably also shed branches faster than slower growing ones. If slower growing trees suffered from higher liana loads they would have been more likely to die than those faster growing ones with lower loads. When the potential for a tree to grow tall was high, the relative costs of branch shedding would be low, i.e. they are ‘affordable’, compared with the converse. An important qualification to the analysis is that the rgr models were based on the growth of final survivors over the 30-yr period, and not on the rgr they attained up until the time they died on a shorter interval-wise basis using the tree census data (cf. Newbery and Ridsdale 2016).

### Tree-liana interactions

Loss of lianas is closely connected with death of their hosts. Tree dynamics found using dates near to the two liana censuses can be more accurately derived by averaging over to the four periods between the main tree censuses of 1986, 1996, 2001, 2007 and 2015 (Newbery et al. 1999, 2011; and unpubl. data). These further estimates reduce the length of interval bias inherent to mortality rate calculations (Sheil and May 1996). The first and last tree censuses were 2-3 years prior to the liana ones, and showed slightly lower differences in rates between plots compared with those based on the liana-censused trees. For all trees, MP1 had a 33-35% higher annualized mortality rate (*m_a_*) than MP2 across three size classes 30 - < 60, 60 -< 120 and ≥ 120 cm gbh (Appendix S1, Table S2). For medium-sized trees, this proportional difference between plots was similar for the Dipterocarpaceae, but it was > 60% for the Euphorbiaceae. Plots differed hardly at all for large and very large dipterocarps (1-8%) but were somewhat lower for large euphorbs (20%). Because of their life-form, species in the Euphorbiaceae hardly ever reach the very large size class to allow estimates.

Between 1986 and 2015 densities of trees (N/ha) ≥ 30 cm gbh changed from 484 to 452 in MP1, and 455 to 477 in MP2. The corresponding basal area abundances (m^2^/ha) were 26.34 to 27.14 in MP1 and 26.79 to 29.28 in MP2. These density shifts account for the differing percent changes in numbers of lianas per tree, when based on number of lianas per plot area and mean number per tree. Over the almost 30 years, MP1 with the higher mortality, particularly of very large trees, decreased in tree density and made little gain in basal area, but MP2 with lower mortality, increased in density and basal area, having more very large trees in MP2 than MP1 (Newbery et al. 1992, and unpubl. data; Appendix S1: Fig. S1). In addition, largest-tree, that is maximum, heights (≥ 100 cm dbh, bar one at 90 cm) were higher in MP2 than MP1 by 8.1 m at the end of 2016 (62.1 vs 54.0 m, *n* = 19 and 23) (D. M. Newbery, unpubl. data). Again, it is the important local plot-scale variation in the forest structure and dynamics that provided detailed insights into tree-liana interactions.

Of the main census trees with gbh ≥ 240 cm in 1986 (*n* = 89), the proportions dying by 2015 were 33% (13/39) and 48% (24/50) in MP1 and MP2 respectively (χ^2^ = 1.94, df = 1, *P* = 0.16), or as *m_a_* over the 28.74 years, 1.40 and 2.25 %/yr. In the liana census data, the number alive at 1988 were 36 and 44 respectively, just one death in MP1 between 1986 and 1988. These *m_a_*-values are indeed the reverse of those for the very large trees (gbh ≥ 120 cm) in respect to plot differences. Nevertheless, the apparent trends among the very largest trees in number of lianas with increasing gbh (Fig. 3, Appendix S1, Fig. S2a, b) is due to three liana counts of 20, 18 (MP1), and 13 (MP2): one tree dying, two surviving. In both plots there were only three trees with ≥ 7 lianas each.

Whilst tree mortality rates were higher in MP1 than MP2 for all three size classes, it was the disproportionately higher-laden trees in the very large size class that led to more lianas being lost in MP1 than in MP2 in that size class. The large and very large trees in MP1 most likely had higher liana loads than those in MP2 even before 1988. In MP1 they had then more lianas to lose by 2018 than those in MP2 by 2018. Without precise data on large branch fall rates, the large to very-large trees must be assumed to have been shedding their lianas at similar rates in MP1 and MP2. Therefore, the difference between the plots, for a majority of the dipterocarps at least, was a result of tree mortality over and above a higher branch fall loss in very large trees compared with large and medium-sized ones.

For trees overall to show similar changes in distributions of numbers of lianas per tree in both plots implies a different process to have been operating in MP2 from that in MP1. The main changes were in the medium-sized understorey trees, where tree mortality was much higher in MP2 than MP1. However, survivors in these size classes were also slowly accumulating lianas, perhaps as newly-rooted ones or coming from fallen trees (Putz 1984). Recruits, being younger and smaller than other trees, would have had little time to accumulate lianas, and hence their loads were relatively low by 2018. The different dynamics in the two plots resulted in similar frequency distributions of lianas per tree though. The overstorey dipterocarps seemingly have more the option of branch shedding to lose lianas, but the understorey euphorbs far less (Campbell and Newbery 1993). The euphorbs can for the most part only lose their lianas on the death of the host. An interesting feature is the higher proportion of trees with no lianas in MP2 than MP1, particularly for the dipterocarps.

If fast-growing medium-sized dipterocarps (Newbery *et al*. 1999), moving from under- to overstorey, were losing branches they might well have shed their lianas along with them. Shedding lianas with branches is one obvious way to relieve the load and recover rgr for onward growth for an overstorey species, and thereby increase its survival under competition with not just the lianas but also neighbouring trees. As understorey species do not grow into the overstorey at all, these considerations are not so important for them. Understorey trees incidentally provide a means for lianas to ‘step up’ onto neighbouring overstorey trees. If large trees remained infested in their canopies and yet have no lianas on their own stems at near-ground level, these lianas are presumably coming across from neighbouring medium-sized trees. This would cause a mismatch between stem counts and liana effects.

The influence of spatial autocorrelation on the fitted values of the independent variables’ coefficients was often quite small, both when the dependent variable was number of lianas per tree (regressed on tree size and elevation) and when it was tree response as growth or survival (regressed on number of lianas per tree and tree size). This indicates that spatial aggregation of numbers of lianas per tree was low and near to random, i.e. locally trees with many lianas and trees with few lianas were well mixed and not forming patches of similarly high or low liana-laden trees in the forest. This form of (spatial) aggregation is not to be confused with the (focal) aggregation of lianas themselves on individual trees (see Campbell and Newbery 1993). If tree growth and survival depended strongly on tree size, and trees in terms of their sizes were randomly distributed in the forest, local neighbourhoods would not have shown correlated growth responses, again supporting a near random distribution of responses spatially. The diffuse spread of lianas in and between crowns would also have diluted any loci of raised or lowered responses. The continual turnover of all size classes of focus trees and their neighbours, due to growth advancement, mortality and recruitment, would moreover add further to the spatial variability as a form of neighbourhood stochasticity (Newbery and Lingenfelder 2009, Newbery and Stoll 2013, Newbery and Ridsdale 2016). The lack of a spatial autocorrelation might suggest additionally that there were in fact very few large forest gaps created in recent decades, ones that had let in enough light to restart patchy secondary succession; and that the effects of thinning in the canopy during ENSO dry periods were broadly diffuse and scattered in terms of the temporarily enhanced light levels reaching the forest understorey.

### Lianas in late succession

Forest successional development was most likely different in MP1 and MP2 up until 1988. To have more lianas by then on the very large trees in MP1 than in MP2 might indicate contingent historical processes, which implies that MP1 was perhaps more open than MP2 in the past. As a consequence, MP1 gained more lianas than MP2, and these were carried on up to 1988. If the thesis that the forest at Danum is now in an end successional phase after a major disturbance ∼120 yrs prior to the first liana census (Newbery et al. 1992) is true, and since then intermittently ENSO events have been occasionally thinning the overstorey through small branch and twig abscission and thereby lighting the understorey (Walsh and Newbery 1999; Newbery et al. 1999, 2011), liana densities and dynamics reflect an asynchrony between the plots in their successional recovery. Once lianas are well established, ENSO events probably provide the means to put on more mass in the crowns of all storeys. The two strong events in this study period were the 1982-83 and 1997-98 ENSOs (Newbery and Lingenfelder 2009), but moderate ones also occurred in 2010-11 and 2015-16 (based extension on the Accumulated Rainfall Anomaly index of Newbery & Lingenfelder 2009: Fig. 1). The first event came 3-4 years before the main plots census in 1986 and was likely influential on the 1988 liana densities. Nunes et al. (2019) have shown that during the last event there was an increase in leaf litterfall at Danum.

This long-term forest process was probably continuing to converge between 1988 and 2018, with plot-level tree basal area moving towards its maximum faster in, or more ahead of, MP2 than MP1. Mature closed forests approaching, or already at, the site-defined maximum carrying capacity (Newbery et al. 1992) might be expected to lessen their liana loads because as the forest became denser in basal area terms, branches that fell would be less easily replaced on the large trees in the overstorey, and lianas would have increasingly less favorable light conditions in which to recruit and grow on the small and medium-sized trees. Heavy liana cover in the understorey would further limit light reaching tree leaves, and hasten their decline under competition with less infested neighbours (Schnitzer and Bongers 2011). By contrast, any large branch fall also perturbs the canopy by causing small openings, and these let in more light to the understorey and hence promote tree and liana growth.

Species with high mean numbers of lianas per tree in 1988 had much larger decreases by 2018 than those with low numbers per tree (e.g. from 3.0 vs 1.0 to ∼1.9 vs 0.9). This indicated a form of negative density dependence at the community level, perhaps by the more heavily laden species disproportionally shedding more lianas or dying from them, or recruiting them more slowly. However, species’ numbers of lianas per tree in 1988, in 2018, and the differences between censuses, were not correlated with mortality (%*d*) or growth rate of stems (rgr). Tree recruitment into the ≥ 30 cm gbh population might have been expected to have been highly correlated with growth rates of small trees (10 -< 50 cm gbh). This suggests that each species was individually determined by different combinations of the factors which led to their gains and losses of lianas.

The increased separation of a group of several of the dipterocarp species in terms of numbers of lianas per tree and OUI, or similarly ln(gbh), between 1988 and 2018 highlighted an interesting dynamic conditional on a strong family trait. Differences in numbers between censuses reflected how the frequency distributions were changing with increasing size class. Even if branch shedding remains the leading hypothesis, other factors are important, e.g. the tendency for medium-sized dipterocarps to have faster stem growth rates than most other species, which is contributory to their upward advancement towards the canopy (Newbery et al. 1999).

The species-level analysis revealed four species which did not belong to the group with low numbers of lianas. *Shorea fallax* had not so low numbers but an intermediate OUI, whilst *S. pauciflora* had a high mean number of lianas despite its high OUI; *V. dulitensis* and *H. nervosa* were intermediate in both numbers of lianas and OUI-values. Furthermore, *S. pauciflora* unlike the other dipterocarps barely changed between censuses. These differences may be explained as follows: *S. fallax* was unusual among the dipterocarps by having a large proportion of medium-sized trees and hence higher than average numbers of lianas per tree for a dipterocarp, and *V. dulitensis* and *H. nervosa*, though having a tendency to be medium-sized trees too, did not reach the maximum sizes of other dipterocarps (hence all three had intermediate OUIs; D. M. Newbery et al., unpubl. data). *Shorea pauciflora* is somewhat special in that its maximum height was only 48.4 m versus 51.0 – 62.2 m (mean 57.2 m) for four others in the separated group (*S. argentifolia*, *S. johorensis*, *S. parvifolia* and *S. pilosa*; D. M. Newbery unpubl. height data), and the corresponding height/dbh ratios were 0.36 vs 0.45 – 0.51 (mean 0.47). This height difference of 8.8 m may then account for the higher load of unshed lianas on *S. pauciflora*. Once again the details of each species’ dynamics, population structure and architecture are all-important. That the graphs of number of lianas per tree versus ln(rgr) were similar to those for number of lianas versus OUI with respect to the dipterocarps reflects the well-established association between fast growth of juveniles and final maximum size in the overstorey.

Another reason why MP1 had more lianas on its very large trees than MP2 in 1988, might have been topography (see Fig. 1 in Lingenfelder and Newbery 2009). MP1 is on the side of a long pronounced ridge with steep slopes and is relatively topographically distinct, whereas MP2, also with an overall similar difference in elevation of 35 – 40 m over the 400 m plot length, is less steep and forms a spur to a higher backing ridge. These subtle differences in topography likely played a role in water availabilities during dry periods, the top of MP2 being less prone to dryness than the ridge top in MP1 (N. Chappell, pers. comm.). Past drought effects could *ex hypothesis* have been stronger in MP1 than MP2, making larger lighter areas in the understorey, and this promoting the establishment of more lianas in the former than latter, with lasting consequences.

Although the number of lianas decreased with elevation there was a similarly strong relationship in both plots: so differences in elevation — at least in recent decades — were not a good explanation for different liana changes across plots. Very large trees were also quite evenly spread across the topographic gradient in both plots (D. M. Newbery, unpubl. data). The proportions of the very largest trees (gbh ≥ 240 cm) dying in the lower and upper halves of the plots (corresponding roughly to lower slopes and ridges) were 35 and 32% in MP1 and 46 and 50% in MP2 (χ^2^ = 0.05, *P* = 0.82; χ^2^ = 0.09, *P* = 0.77). So host size distribution was not an explanation either. This would then indicate that MP2 probably recovered faster than MP1 in successional terms and any very large trees that did have high loads in MP2 had already fallen before 1988, and in the 1988-2018 period a similar phase was being realized in MP1. Meanwhile MP2 was showing a next, maybe final, phase of liana loss within the understorey. Indeed, if intensity of disturbance was less in MP2 than MP1, a faster recovery (less impeded by lianas) would have been expected more in the former than the latter. This represents a shifting mosaic in time and space, in theory reminiscent of the ‘pattern and process’ idea of Watt (1947), applied by Richards (1952) to rain forests. The dynamics of lianas in such a late successional forest can be linked historically to liana activity soon after that distant strong disturbance set the long-term regrowth on its variable course.

### A new hypothesis for canopy roughness

If during succession and forest maturity liana loads reach a quasi-stable equilibrium density and biomass (Muller-Landau and Visser 2019, Muller-Landau and Pacala 2020), this would be expected in mid-succession when there remained sufficient light for recruitment and onward liana spread; and then follows decline as the forest canopy fully closed. By how much lianas might decline could depend on the time until a next major disturbance. If lianas established well on understorey trees, they would respond positively to the ENSO dry periods and at the same time provide more cover for saplings and small trees of the canopy species during these perturbations, which reinforces the over-understorey feedback model of Newbery et al. (1999). The ENSO ‘pushes’ would offset the late successional closing of the canopy (Newbery et al. 2011). As large trees matured and moved from the canopy into emergent storey, they may shed larger branches with heavy liana loads, or lianas are pulled away by being connected to neighbouring trees that fall. This would free their crowns of light competition by lianas, and in dropping branches leave an increasing length of branch-free bole along which it would be difficult for lianas coming from lower crowns to cross onto and re-infest. Essentially, they could ‘break free’ and ‘escape’. Large branch fall would, furthermore, drag connecting lianas to neighbours with them and thereby lessen the chance of re-infestation. The escape hypothesis provides an interesting explanation why this region of NE Borneo, relatively free of storms and below the cyclone belt, has so many standing dead emergent trees (Gale and Hall 2001, Lingenfelder and Newbery 2009, D. M. Newbery pers. obs.), and as a consequence why the canopies of forests of central Sabah are so obviously uneven (Newbery et al. 1992, 1999; D.M.N. reconnaissance flight. By contrast, smaller understorey trees would not have been able to shed their lianas and continued to accumulate them over time until they died with the tree, a further burden for suppressed individuals. Being liana free might also contribute to the stability of exceptionally tall dipterocarps recently recorded for Sabah (Jackson et al. 2021).

The liana shedding hypothesis would challenge Ashton and Hall’s (1992) drought effects one for canopy roughness of Bornean lowland dipterocarp forests, but it could also be seen to complement it if large trees lost large branches more frequently in dry periods than in wet ones. Whilst lianas have the advantage of better growth in dry periods (Schnitzer 2005, 2018), for very large trees that response would eventually lead to their demise when their hosts shed them along with branches. A within-crown feedback process would down-regulate liana abundances in a density dependent manner. It is one mechanism that might operate in the host-liana (-parasite) model of Muller-Landau and Pacala (2020). Since lianas do have strong competitive and growth-suppressing effects on trees, then species with the most successful survivors at the site can be expected to have evolved mechanisms to free themselves of their parasites. The cost of losing lower branches, measured in terms of temporarily reduced photosynthetic activity, should be less than the effect of sustaining a suppressing liana load that could limit the same longer-term. Losing branches would also allow more light reaching juvenile trees in the understorey which, in the Dipterocarpaceae especially, tend to be clustered around their adults at Danum (Stoll and Newbery 2005). Pattern and extent of branch-fall calls for more detailed field recording and a clearer interpretation within a framework of tree architecture, branch growth and survival (Rutishauser et al. 2011).

The liana shedding hypothesis places emphasis on how a host tree can minimize liana loads and improve its fitness, whereas the commoner line in the literature has been more about how the lianas can maximize their hold on hosts and their fitness. Ultimately this would be played out in their respective life-history schedules. However, to properly model equilibrium dynamics, the inherently stochastic long- and short-term nature of the environment of N.E. Borneo — in the form of recoveries from past major disturbances and continuing ENSO perturbations, needs to be incorporated.

## Acknowledgements

We thank R. Ewers of Imperial College London for arrangements for the MSc work of C.Z. at Danum, R. Nilus (Sabah Forest Department) for local collaboration, A. F. Karolus for field assistance (Danum Valley Field Centre), and M. Lingenfelder (Bern University) for the Danum tree database management and taxonomic codes matching. Permission for C.Z. to conduct the field work was granted by the Danum Valley Management Committee and the Sabah Biodiversity Council. Two anonymous reviewers are thanked for their improvements to the paper.

## Author contributions

D.M.N. designed the set-up at Danum and conceived the study. C.Z. undertook the 2018 liana field census, the first analyses, and prepared the R-code for the randomization procedure. D.M.N. extended the statistical analysis and interpretation, and largely wrote the paper.

## Funding sources

The first liana census in 1988, conducted by E. J. F. Campbell, was an extension to tree plot work made with a grant from the Natural Research Council in UK (GR3/5555), and the analysis of the second liana census data drew on results of the main plot tree censuses to 2007 funded more recently by the Swiss National Science Foundation (Grants 3100-59088 and 31003A-110250). The last plot census in 2015 for the trees was financially supported by Chair for Vegetation Ecology (D.M.N.) at Bern.

### Data Accessibility

The data from the two liana censuses are available at the Dryad Digital Repository: (DOI: 10.5061/dryad.z8w9ghx8t).

