## Supplementary Materials for "Change in liana density over 30 years in a Bornean rain forest supports the escape hypothesis"

**Appendix S1. Supporting results**

Table S1. Number of trees, relative abundance, proportion of liana infested trees and average number of lianas per tree for all trees and those in the third to tenth most abundant families, in each census in main plots (a) MP1 and (b) MP2. These data extend those in Table 1 of the main text.

| (a) MP1 | Number of trees in plot | | Trees with lianas (%) | | Average number of lianas per tree | |
|---|---|---|---|---|---|---|
| Tree family | 1988 | 2018 | 1988 | 2018 | 1988 | 2018 |
| Meliaceae | 144 | 134 | 59.0 | 70.9 | 2.40 | 1.94 |
| Phyllanthaceae | 130 | 113 | 74.6 | 81.4 | 2.72 | 2.16 |
| Lauraceae | 126 | 143 | 54.8 | 71.3 | 1.67 | 1.66 |
| Annonaceae | 116 | 102 | 50.9 | 65.7 | 1.50 | 1.58 |
| Myrtaceae | 100 | 95 | 62.0 | 67.4 | 2.00 | 1.86 |
| Malvaceae | 98 | 104 | 59.2 | 68.3 | 2.16 | 1.80 |
| Sapotaceae | 89 | 92 | 74.2 | 83.7 | 2.70 | 2.32 |
| Fagaceae | 83 | 66 | 56.6 | 56.1 | 2.46 | 1.44 |

| (b) MP2 | Number trees in plot | | Trees with lianas (%) | | Average number of lianas per tree | |
|---|---|---|---|---|---|---|
| Tree family | 1988 | 2018 | 1988 | 2018 | 1988 | 2018 |
| Meliaceae | 105 | 142 | 61.9 | 67.6 | 2.39 | 1.73 |
| Phyllanthaceae | 104 | 104 | 74.0 | 69.2 | 2.84 | 1.63 |
| Lauraceae | 94 | 101 | 63.8 | 61.4 | 1.93 | 1.50 |
| Annonaceae | 98 | 113 | 69.4 | 56.6 | 2.39 | 1.37 |
| Myrtaceae | 86 | 71 | 73.3 | 66.2 | 3.40 | 1.67 |
| Malvaceae | 79 | 80 | 50.6 | 66.3 | 1.86 | 1.65 |
| Sapotaceae | 41 | 51 | 63.4 | 74.5 | 2.78 | 2.18 |
| Fagaceae | 66 | 64 | 39.4 | 57.8 | 1.85 | 1.41 |

Table S2. Average annualized mortality rates ($m_a$) over the four periods between censuses at Danum in the two plots, MP1 and MP2 (1986-96, 1996-2001, 2001-07, 2007-15), for trees ≥ 30 cm gbh at the starts of intervals, in three size classes (scl: 1, 30-< 60; 2, 60 -< 120; and ≥ 120 cm gbh), found for all trees in all families (All), those in the Dipterocarpaceae (Dipt) and those in the Euphorbiaceae (Euph), with proportional differences (Diff.) in rates for MP1 over MP2.

| | | $m_a$ (%/yr) | | Diff.[a] |
|---|---|---|---|---|
| fam | scl | MP1 | MP2 | (%) |
| All | 1 | 2.50 | 1.88 | 33.0 |
| | 2 | 2.41 | 1.81 | 33.2 |
| | 3 | 2.07 | 1.53 | 35.3 |
| Dipt | 1 | 2.70 | 1.99 | 35.7 |
| | 2 | 1.86 | 1.72 | 8.1 |
| | 3 | 1.61 | 1.62 | −0.6 |
| Euph[b] | 1 | 3.67 | 2.27 | 61.7 |
| | 2 | 3.41 | 2.83 | 20.5 |
| | 3 | -- | -- | -- |

[a] as $(m_{a\text{-MP1}} - m_{a\text{-MP2}}) / m_{a\text{-MP2}}$; [b] for scl = 3, too few trees to estimate $m_a$.

Fig. S1. Ln-ln plots of numbers of trees ≥ 30 cm gbh (nr_tree) versus mean size class gbh in the two main plots (a) MP1, and (b) MP2, at Danum, for the two liana census dates 1988 and 2018. Slopes of the lines (± SE) from linear regression [and the adjusted $R^2$-values of the fits] were: MP1, 1988, −1.959 ± 0.188 [82.5%]; 2018, −1.940 ± 0.160 [86.4%]; MP2, 1988, −1.967 ± 0.151 [88.0%]; 2018, −1.869 ± 0.117 [91.7%].

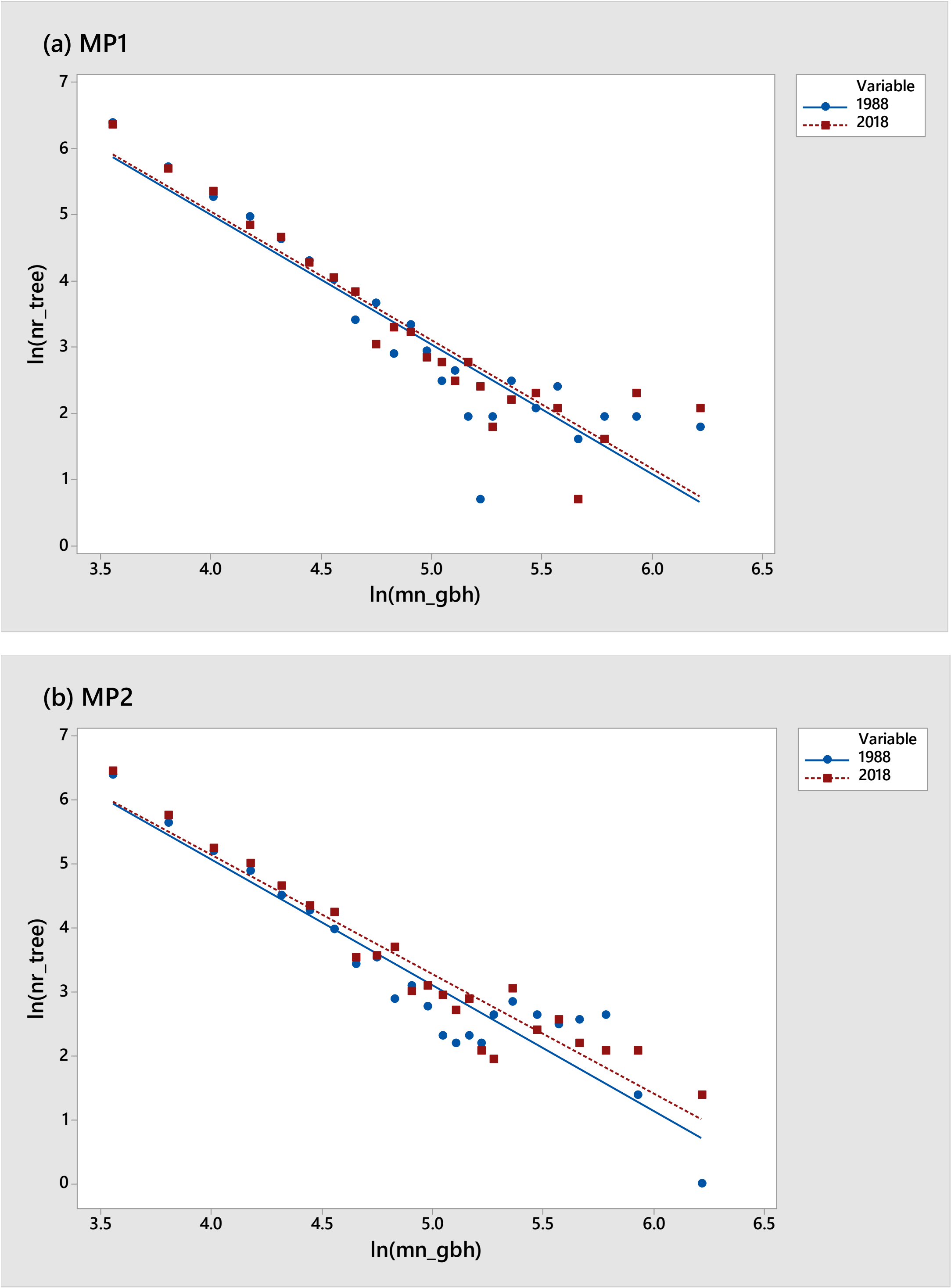

Fig. S2. Relationships from negative binomial GLM regressions for the number of lianas per tree (nr_liana) versus tree size (gbh – ln-transformed) in main plots 1 and 2, at each census date: solid line, 1988; dashed line, 2018, as simplified in Fig. 3 main text, but here showing the scatter of values for the individual trees.

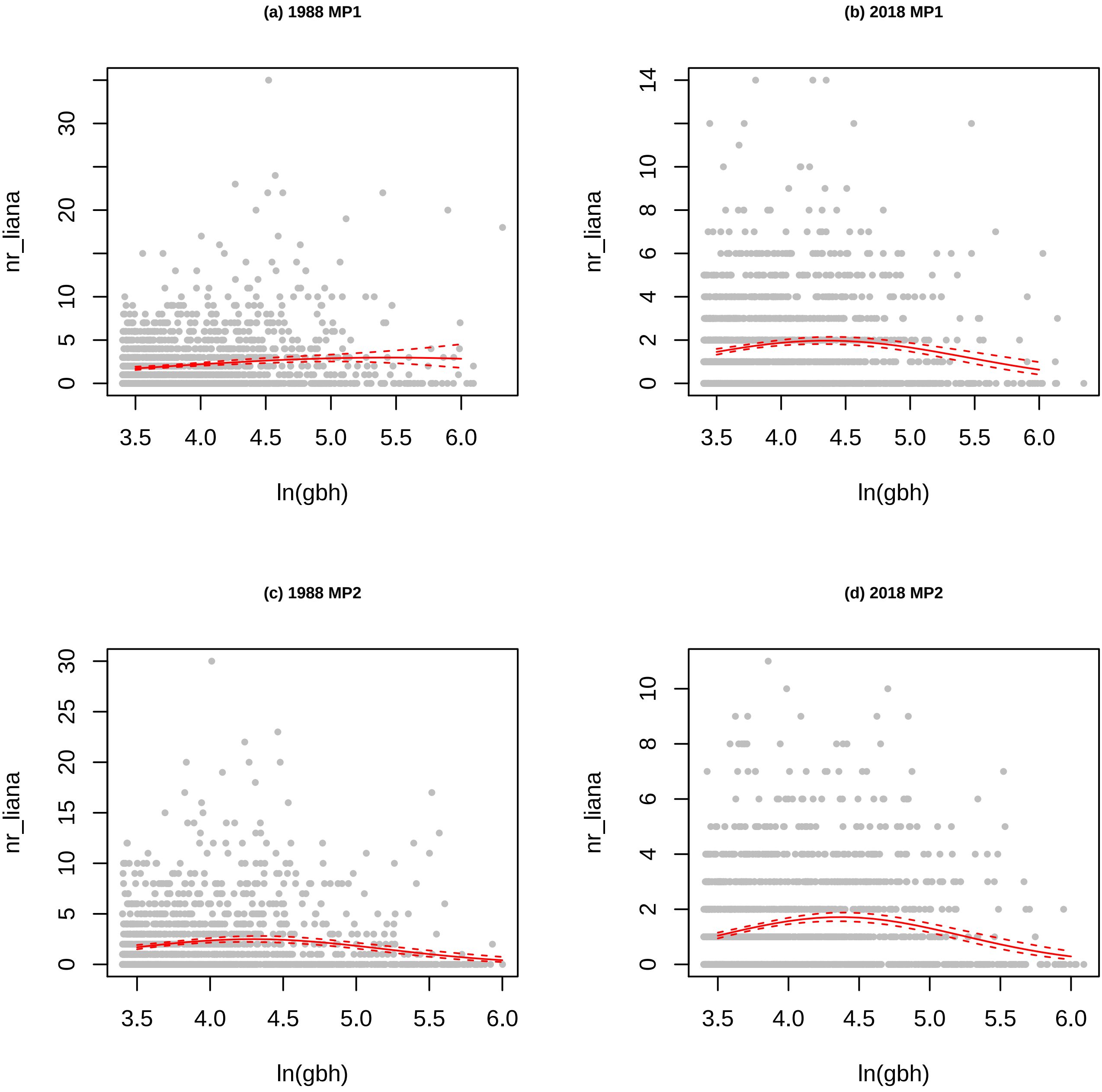
(a) 1988 MP1
(b) 2018 MP1
(c) 1988 MP2
(d) 2018 MP2
nr_liana
ln(gbh)
0
5
10
20
30
2
4
6
8
12
14
15
25
3.5
4.0
4.5
5.0
5.5
6.0

### Appendix S2. Regression model statistics

Probability levels: ***, $P \leq 0.001$; **, $P \leq 0.01$; *, $P \leq 0.05$; °, $P \leq 0.10$, $^{ns}$, $P > 0.10$.

Table S1a. Regressions of numbers of lianas per tree on tree girth at breast height (gbh) and elevation (GLM, negative binomial error) in the censuses of 1988 and 2018.

| | | MP1 | | MP2 | |
|---|---|---|---|---|---|
| census | term | est ± se | *t* | est ± se | *t* |
| 1988 | intercept | −2.891 ± 1.634 | −1.769$^{ns}$ | −9.989 ± 1.981 | −5.041** |
| | ln(gbh) | 1.533 ± 0.757 | 2.026* | 5.157 ± 0.925 | 5.572*** |
| | [ln(gbh)]$^2$ | −0.1385 ± 0.0864 | −1.602$^{ns}$ | −0.6006 ± 0.1064 | −5.643*** |
| | elevation | −0.01467 ± 0.00428 | −3.430*** | −0.00939 ± 0.00334 | −2.810** |
| 2018 | intercept | −6.740 ± 1.434 | −4.701*** | −11.873 ± 1.613 | −7.361*** |
| | ln(gbh) | 3.494 ± 0.670 | 5.214*** | 5.812 ± 0.758 | 7.671*** |
| | [ln(gbh)]$^2$ | −0.4014 ± 0.0774 | −5.183*** | −0.6650 ± 0.0878 | −7.576*** |
| | elevation | −0.01162 ± 0.00356 | −3.269** | −0.01605 ± 0.00255 | −6.296*** |

Table S1b. Regressions of numbers of lianas per tree on tree girth at breast height (gbh) and elevation (HGLM, quasi-Poisson error) in the censuses of 1988 and 2018.

| | | MP1 | | MP2 | |
|---|---|---|---|---|---|
| census | term | est ± se | *t* | est ± se | *t* |
| 1988 | intercept | −2.760 ± 1.452 | −1.902° | −9.352 ± 1.866 | −5.013*** |
| | ln(gbh) | 1.493 ± 0.671 | 2.226* | 4.841 ± 0.878 | 5.512*** |
| | [ln(gbh)]$^2$ | −0.1428 ± 0.0765 | −1.866° | −0.5711 ± 0.1021 | −5.596*** |
| | elevation | −0.01478 ± 0.00391 | −3.783*** | −0.01000 ± 0.00294 | −3.402** |
| 2018 | intercept | −6.769 ± 1.480 | −4.575*** | −11.504 ± 1.620 | −7.102*** |
| | ln(gbh) | 3.511 ± 0.695 | 5.052*** | 5.626 ± 0.763 | 7.370*** |
| | [ln(gbh)]$^2$ | −0.4057 ± 0.0808 | −5.018*** | −0.6428 ± 0.0889 | −7.231*** |
| | elevation | −0.01093 ± 0.00347 | −3.148** | −0.01635 ± 0.00242 | −6.750*** |

Table S1c. Regressions of numbers of lianas per tree on tree girth at breast height (gbh) and elevation (GLM, negative binomial error) in the censuses of 1988 and 2018 for trees in the Euphorbiaceae(Euph) and Dipterocarpaceae (Dipt) respectively. The significance levels in parenthesis are the result of HGLM accounting spatial autocorrelation.

| | MP1-1988, Euph | | MP2-2018, Dipt | |
|---|---|---|---|---|
| term | est ± se | *t* | est ± se | *t* |
| intercept | −24.983 ± 9.394 | −2.659** [*] | −12.803 ± 5.268 | −2.431* [*] |
| ln(gbh) | 12.926 ± 4.754 | 2.719** [*] | 6.237 ± 2.343 | 2.663** [**] |
| [ln(gbh)]$^2$ | −1.5820 ± 0.5958 | −2.655** [*] | −0.7004 ± 0.2554 | −2.742** [**] |
| elevation | −0.02383 ± 0.00987 | −2.415* [*] | −0.02271 ± 0.01046 | −2.172* [$^{ns}$] |

Table S2a. Logistic regressions of tree survival 1988-2018 on number of lianas per tree and tree girth at breast height (gbh) in 1988, and elevation (GLM, binomial error).

| | MP1 | | MP2 | |
|---|---|---|---|---|
| term | est ± se | *t* | est ± se | *t* |
| intercept | −0.776 ± 0.379 | −2.047* | −0.370 ± 0.383 | $-0.966^{ns}$ |
| $nlianas^{1/2}$ | −0.1449 ± 0.0482 | −3.007** | −0.2008 ± 0.0482 | −4.169*** |
| ln(gbh) | 0.1948 ± 0.0913 | 2.133* | 0.1530 ± 0.0901 | $1.697^{o}$ |
| elevation | −0.00174 ± 0.00607 | $-0.287^{ns}$ | 0.01210 ± 0.00436 | 2.772** |

Table S2b. Regressions of tree survival 1988-2018 on number of lianas per tree and tree girth at breast height (gbh) in 1988, and elevation (GLM, binomial error), under repeated random sampling per subplot. Statistics are means of values from $N'$ = 500 runs: each regression had $n$ = 100 trees. All $P(t) > 0.05$.

| | MP1 | | MP2 | |
|---|---|---|---|---|
| term | est ± se | *t* | est ± se | *t* |
| intercept | −0.8445 ± 1.6567 | −0.504 | −0.6336 ± 1.6405 | −0.357 |
| $nlianas^{1/2}$ | −0.1125 ± 0.2056 | −0.533 | −0.1826 ± 0.2025 | −0.902 |
| ln(gbh) | 0.1949 ± 0.3995 | 0.486 | 0.2186 ± 0.3851 | 0.527 |
| elevation | −0.00060 ± 0.02604 | −0.021 | 0.01217 ± 0.01825 | 0.657 |

Table S2c. GLMM regressions of tree survival 1988-2018 on number of lianas per tree and tree girth at breast height (gbh) in 1988, and elevation (binomial error), accounting for spatial autocorrelation (cluster = subplot), for trees in the Dipterocarpaceae.

| | MP1 | | MP2 | |
|---|---|---|---|---|
| term | est ± se | *t* | est ± se | *t* |
| intercept | 0.652 ± 0.832 | $0.784^{ns}$ | 0.926 ± 0.897 | $1.033^{ns}$ |
| $nlianas^{1/2}$ | −0.2020 ± 0.1295 | $-1.640^{ns}$ | −0.4064 ± 0.1327 | −3.064** |
| ln(gbh) | 0.0999 ± 0.1839 | $0.543^{ns}$ | −0.0781 ± 0.1746 | $-0.447^{ns}$ |
| elevation | −0.03859 ± 0.01983 | $-1.946^{o}$ | 0.00482 ± 0.01264 | $0.381^{ns}$ |

Table S2d. GLMM regressions of tree survival 1988-2018 on number of lianas per tree and tree girth at breast height (gbh) in 1988, and elevation (binomial error), accounting for spatial autocorrelation (cluster = subplot), for trees in the Euphorbiaceae.

| | MP1 | | MP2 | |
|---|---|---|---|---|
| term | est ± se | *t* | est ± se | *t* |
| intercept | −2.147 ± 1.925 | $-1.115^{ns}$ | 0.596 ± 1.823 | $0.327^{ns}$ |
| $nlianas^{1/2}$ | −0.4311 ± 0.1898 | −2.272* | −0.3445 ± 0.1380 | −2.497* |
| ln(gbh) | 0.3864 ± 0.4946 | $0.781^{ns}$ | −0.1827 ± 0.4801 | $-0.381^{ns}$ |
| elevation | −0.00411 ± 0.01986 | $0.207^{ns}$ | 0.01513 ± 0.01298 | $1.165^{ns}$ |

Table S3a. Linear regressions of tree relative growth rate (rgr) 1988-2018 on number of lianas per tree and tree girth at breast height (gbh) in 1988 (LM, gaussian error).

| | MP1 | | MP2 | |
|---|---|---|---|---|
| term | est ± se | *t* | est ± se | *t* |
| intercept | 0.781 ± 0.071 | 11.067*** | 0.610 ± 0.059 | 10.262*** |
| nlianas$^{1/2}$ | −0.0364 ± 0.0095 | −3.838*** | −0.0328 ± 0.0082 | −3.979*** |
| ln(gbh) | −0.1080 ± 0.0171 | −6.309*** | −0.0791 ± 0.0143 | −5.523*** |

Table S3b. Spatial autoregressions of tree relative growth rate (rgr) 1988-2018 on number of lianas per tree and tree girth at breast height (gbh) in 1988 (GLS, gaussian error), based on a correlogram, and using an inferred distance matrix with neighbour range 0-20 m).

| | MP1 | | MP2 | |
|---|---|---|---|---|
| term | est ± se | *t* | est ± se | *t* |
| intercept | 0.759 ± 0.071 | 10.727*** | 0.597 ± 0.060 | 10.011*** |
| nlianas$^{1/2}$ | −0.0371 ± 0.0095 | −3.921*** | −0.0347 ± 0.0082 | −4.203*** |
| ln(gbh) | −0.1030 ± 0.0171 | −6.0358*** | −0.0747 ± 0.0143 | −5.238*** |

Table S3c. Linear regressions of tree relative growth rate (rgr) 1988-2018 on number of lianas per tree and tree girth at breast height (gbh) in 1988 (LM, gaussian error), under repeated random sampling per subplot. Statistics are means of values from *N*' = 500 runs: each regression had *n* = 100 trees.

| | MP1[a] | | MP2 | |
|---|---|---|---|---|
| term | est ± se | *t* | est ± se | *t* |
| intercept | 0.790 ± 0.205 | 3.855*** | 0.620 ± 0.187 | 3.311*** |
| nlianas$^{1/2}$ | −0.0377 ± 0.0257 | −1.469$^{ns}$ | −0.0299 ± 0.0256 | −1.168$^{ns}$ |
| ln(gbh) | −0.1088 ± 0.0498 | −2.185* | −0.0800 ± 0.0449 | −1.780$^{o}$ |

Table S3d. General least squares regressions of tree relative growth rate (rgr) 1988-2018 on number of lianas per tree and tree girth at breast height (gbh) in 1988 (GLS, mixed-model, gaussian error), accounting for spatial autocorrelation (corExp variogram), for trees in the Dipterocarpaceae.

| | MP1 | | MP2 | |
|---|---|---|---|---|
| term | est ± se | *t* | est ± se | *t* |
| intercept | 1.744 ± 0.149 | 11.682*** | 1.435 ± 0.145 | 9.901*** |
| nlianas$^{1/2}$ | −0.0320 ± 0.0233 | −1.373$^{ns}$ | −0.0330 ± 0.0245 | −1.350$^{ns}$ |
| ln(gbh) | −0.2736 ± 0.0320 | −8.588*** | −0.2153 ± 0.0297 | −7.262*** |

Table S3e. General least squares regressions of tree relative growth rate (rgr) 1988-2018 as for Table S3d, accounting for spatial autocorrelation (corExp variogram), for trees in the Euphorbiaceae.

| | MP1 | | MP2 | |
|---|---|---|---|---|
| term | est ± se | *t* | est ± se | *t* |
| intercept | 0.262 ± 0.198 | 1.324$^{ns}$ | 0.288 ± 0.122 | 2.352* |
| nlianas$^{1/2}$ | −0.0034 ± 0.0203 | −0.166$^{ns}$ | −0.0230 ± 0.0104 | −2.217* |
| ln(gbh) | −0.0228 ± 0.0511 | −0.445$^{ns}$ | −0.0368 ± 0.0325 | −1.133$^{ns}$ |

Table S4. Regressions of numbers of lianas per tree in 1988 and 2018, on tree relative growth rate 1988-2018 (rgr) and girth at breast height (gbh) for the census, and elevation (GLM, negative binomial error).

| | | MP1 | | MP2 | |
|---|---|---|---|---|---|
| census | term | est ± se | *t* | est ± se | *t* |
| 1988 | intercept | −1.320 ± 2.447 | $-0.539^{ns}$ | −10.890 ± 2.772 | −3.928*** |
| | $rgr_{88\text{-}18}$ | −0.4856 ± 0.2041 | −2.380* | −0.8052 ± 0.2213 | −3.638*** |
| | $\ln(gbh_{88})$ | 1.029 ± 1.126 | $0.914^{ns}$ | 5.597 ± 1.297 | 4.315*** |
| | $[\ln(gbh_{88})]^2$ | −0.1035 ± 0.1275 | $-0.812^{ns}$ | −0.6556 ± 0.1492 | −4.392*** |
| | elevation | −0.01875 ± 0.00673 | −2.787** | −0.00536 ± 0.00483 | $-1.109^{ns}$ |
| 2018 | intercept | −7.880 ± 2.287 | −3.446*** | −19.058 ± 2.450 | −7.780*** |
| | $rgr_{88\text{-}18}$ | −0.6961 ± 0.1699 | −4.098*** | −0.8944 ± 0.1799 | −4.972*** |
| | $\ln(gbh_{18})$ | 4.167 ± 1.022 | 4.078*** | 9.011 ± 1.108 | 8.136*** |
| | $[\ln(gbh_{18})]^2$ | −0.4765 ± 0.1121 | −4.429*** | −1.0024 ± 0.1233 | −8.129*** |
| | elevation | −0.01514 ± 0.00513 | −2.948** | −0.01696 ± 0.00361 | −4.702*** |

Table S5. On the interaction term in negative binomial GLM models

Model 4: $\text{nlianas} = \beta_0 + \beta_1\cdot\text{date} + \beta_2\cdot\ln(\text{gbh}) + \beta_3\cdot\text{date:}\ln(\text{gbh}) + \beta_4\cdot[\ln(\text{gbh})]^2 + \beta_5\cdot\text{date:}[\ln(\text{gbh})]^2$

From the results in Table 9 (main text), applying: $\beta_{int} = \beta_1 + \beta_3\cdot\ln(\text{gbh}_i) + \beta_5\cdot[\ln(\text{gbh}_i)]^2$:

(a) MP1:

| ln(gbh) | $\beta_{int}$ | $\exp(\beta_{int})$ |
|---|---|---|
| 3.5 | −0.164 | 0.849 |
| 3.75 | *−0.149* | *0.862* |
| *4.0* | *−0.164* | *0.849* |
| 4.5 | −0.288 | 0.750 |
| 5.5 | −0.908 | 0.403 |

(b) MP2:

| ln(gbh) | $\beta_{int}$ | $\exp(\beta_{int})$ |
|---|---|---|
| 3.5 | −0.486 | 0.615 |
| 3.75 | *−0.444* | *0.641* |
| *4.0* | *−0.417* | *0.659* |
| 4.5 | −0.356 | 0.701 |
| 5.5 | −0.320 | 0.726 |

Table S6. Inferred relationships for nlianas vs ln(gbh) in MP1 and MP2, (a) inferred from a GLM model with year as factor included, and (b) in comparison to earlier GLMs for the years separately.

(a) Inferred coefficients from model 4 in Table 9 (main text):

| | Ref “88” | For -> “18” |
|---|---|---|
| MP1 | intercept −3.24263 | + (−3.9479) = −7.19053 |
| | ln(gbh) 1.59732 | + 2.0145 = 3.61182 |
| | $[\ln(gbh)]^2$ −0.14719 | + (−0.26723) = −0.4142 |
| MP2 | intercept −10.40812 | + (−1.83716) = −12.2453 |
| | ln(gbh) 5.27499 | + 0.58736 = 5.86236 |
| | $[\ln(gbh)]^2$ −0.61432 | + (−0.05763) = −0.6720 |

(b) Separate GLM regressions for each year:

| | MP1 | | MP2 | |
|---|---|---|---|---|
| | 1988 | 2018 | 1988 | 2018 |
| intercept | −3.2129 | −7.2876 | −10.436 | −12.1619 |
| ln(gbh) | 1.5835 | 3.6602 | 5.2886 | 5.8200 |
| $[\ln(gbh)]^2$ | −0.14563 | −0.42035 | −0.6160 | −0.6667 |

Table S7. Mean number of lianas per tree as predicted by models 1 to 4, for MP1 and MP2 separately, for all trees and those in the Dipterocarpaceae and Euphorbiaceae. The values are averages of *N*’ = 500 randomizations. Predictions were made with the means of ln(gbh) and $[\ln(gbh)]^2$ taken for plots (a), combined, and (b), separate.

| (a) | | MP1, est ± se | | MP2, est ± se | |
|---|---|---|---|---|---|
| Family | M | 1988 | 2018 | 1988 | 2018 |
| All | 1 | 2.163 ± 0.096 | 1.686 ± 0.078 | 2.017 ± 0.099 | 1.335 ± 0.066 |
| | 2 | 2.141 ± 0.095 | 1.687 ± 0.077 | 2.017 ± 0.099 | 1.335 ± 0.066 |
| | 3 | 2.132 ± 0.094 | 1.685 ± 0.077 | 2.015 ± 0.099 | 1.333 ± 0.066 |
| | 4 | 2.124 ± 0.093 | 1.661 ± 0.076 | 1.960 ± 0.095 | 1.295 ± 0.064 |
| Dipt | 1 | 1.834 ± 0.294 | 0.998 ± 0.170 | 1.364 ± 0.211 | 0.841 ± 0.132 |
| Euph | 1 | 1.918 ± 0.223 | 1.882 ± 0.228 | 1.998 ± 0.273 | 1.045 ± 0.155 |

| (b) | | MP1, est ± se | | MP2, est ± se | |
|---|---|---|---|---|---|
| Family | M | 1988 | 2018 | 1988 | 2018 |
| All | 1 | same as in (a) | -- | -- | -- |
| | 2 | 2.143 ± 0.095 | 1.685 ± 0.077 | 2.017 ± 0.099 | 1.334 ± 0.066 |
| | 3 | 2.136 ± 0.094 | 1.685 ± 0.077 | 2.015 ± 0.099 | 1.333 ± 0.066 |
| | 4 | 2.127 ± 0.093 | 1.663 ± 0.076 | 1.965 ± 0.096 | 1.291 ± 0.064 |
| Dipt | 1 | same as in (a) | -- | -- | -- |
| Euph | 1 | same as in (a) | -- | -- | -- |

### Appendix S3. Comparison of four models of change in lianas

Table S1. Means and standard errors of the fitting statistics for the models 1 – 4 (see main text and Table 9).

(a) MP1

| Variable | Mean | SE Mean | Minimum | Median | Maximum |
|---|---|---|---|---|---|
| AIC.1 | 6365.4 | 4.35 | 6099.0 | 6366.2 | 6650.4 |
| AIC.2 | 6357.9 | 4.31 | 6092.4 | 6357.9 | 6640.2 |
| AIC.3 | 6353.7 | 4.29 | 6087.7 | 6352.5 | 6637.2 |
| AIC.4 | 6341.1 | 4.30 | 6076.3 | 6340.7 | 6633.0 |
| pseudoR2.1 | 0.8670 | 0.0137 | 0.2031 | 0.8585 | 1.9315 |
| pseudoR2.2 | 1.3936 | 0.0193 | 0.4440 | 1.3659 | 2.7016 |
| pseudoR2.3 | 1.7345 | 0.0240 | 0.5610 | 1.6994 | 3.3372 |
| pseudoR2.4 | 2.6418 | 0.0284 | 1.2031 | 2.5737 | 5.0361 |
| dev.1 | 1796.9 | 1.11 | 1726.9 | 1796.6 | 1864.0 |
| dev.2 | 1798.1 | 1.11 | 1729.9 | 1797.8 | 1865.1 |
| dev.3 | 1798.8 | 1.11 | 1731.3 | 1798.6 | 1865.6 |
| dev.4 | 1800.2 | 1.11 | 1732.0 | 1800.0 | 1868.4 |
| df.1 | 1698.3 | 0.956 | 1636.0 | 1697.0 | 1758.0 |
| df.2 | 1697.3 | 0.956 | 1635.0 | 1696.0 | 1757.0 |
| df.3 | 1696.3 | 0.956 | 1634.0 | 1695.0 | 1756.0 |
| df.4 | 1694.3 | 0.956 | 1632.0 | 1693.0 | 1754.0 |
| ano.12.p | 0.02262 | 0.00277 | 0.00000 | 0.00261 | 0.63825 |
| ano.13.p | 0.01191 | 0.00164 | 0.00000 | 0.00055 | 0.42979 |
| ano.14.p | 0.000277 | 0.000085 | 0.000000 | 0.000003 | 0.031996 |
| ano.23.p | 0.05927 | 0.00489 | 0.00001 | 0.01764 | 0.97359 |
| ano.24.p | 0.001937 | 0.000347 | 0.000000 | 0.000071 | 0.104181 |
| ano.34.p | 0.005916 | 0.000968 | 0.000000 | 0.000477 | 0.329315 |
| ano.12.LR | 9.532 | 0.218 | 0.221 | 9.061 | 27.566 |
| ano.13.LR | 15.725 | 0.308 | 1.689 | 15.007 | 40.145 |
| ano.14.LR | 32.285 | 0.395 | 10.559 | 31.308 | 65.398 |
| ano.23.LR | 6.193 | 0.163 | 0.001 | 5.631 | 20.342 |

| | | | | | |
|---|---|---|---|---|---|
| ano.24.LR | 22.753 | 0.353 | 6.158 | 21.831 | 58.006 |
| ano.34.LR | 16.560 | 0.313 | 2.221 | 15.298 | 43.995 |

(b) MP2

| Variable | Mean | SE Mean | Minimum | Median | Maximum |
|---|---|---|---|---|---|
| AIC.1 | 6060.2 | 4.30 | 5764.5 | 6062.0 | 6349.0 |
| AIC.2 | 6061.6 | 4.29 | 5766.5 | 6063.9 | 6350.8 |
| AIC.3 | 6062.2 | 4.30 | 5768.5 | 6064.2 | 6351.1 |
| AIC.4 | 6021.8 | 4.29 | 5720.5 | 6023.2 | 6310.3 |
| pseudoR2.1 | 1.9824 | 0.0218 | 0.7837 | 1.9496 | 3.9091 |
| pseudoR2.2 | 2.0151 | 0.0221 | 0.7863 | 1.9792 | 3.9207 |
| pseudoR2.3 | 2.0934 | 0.0216 | 0.9085 | 2.0522 | 4.0485 |
| pseudoR2.4 | 4.5676 | 0.0334 | 2.7187 | 4.5068 | 7.9302 |
| dev.1 | 1752.8 | 1.08 | 1663.5 | 1753.9 | 1826.4 |
| dev.2 | 1752.8 | 1.08 | 1663.5 | 1753.8 | 1826.3 |
| dev.3 | 1752.9 | 1.08 | 1663.5 | 1753.8 | 1826.2 |
| dev.4 | 1754.6 | 1.08 | 1665.9 | 1755.3 | 1827.7 |
| df.1 | 1744.3 | 0.904 | 1683.0 | 1745.0 | 1806.0 |
| df.2 | 1743.3 | 0.904 | 1682.0 | 1744.0 | 1805.0 |
| df.3 | 1742.3 | 0.904 | 1681.0 | 1743.0 | 1804.0 |
| df.4 | 1740.3 | 0.904 | 1679.0 | 1741.0 | 1802.0 |
| ano.12.p | 0.5737 | 0.0113 | 0.0158 | 0.5849 | 0.9988 |
| ano.13.p | 0.4895 | 0.0122 | 0.0009 | 0.4979 | 0.9995 |
| ano.14.p | 0.000001 | 0.000000 | 0.000000 | 0.000000 | 0.000065 |
| ano.23.p | 0.4282 | 0.0132 | 0.0003 | 0.3906 | 0.9979 |
| ano.24.p | 0.000000 | 0.000000 | 0.000000 | 0.000000 | 0.000021 |
| ano.34.p | 0.000000 | 0.000000 | 0.000000 | 0.000000 | 0.000011 |
| ano.12.LR | 0.5821 | 0.0347 | 0.0000 | 0.2984 | 5.8279 |
| ano.13.LR | 1.9707 | 0.0829 | 0.0010 | 1.3945 | 14.0560 |
| ano.14.LR | 46.387 | 0.477 | 24.443 | 45.558 | 94.141 |
| ano.23.LR | 1.3886 | 0.0789 | 0.0000 | 0.7372 | 13.0457 |
| ano.24.LR | 45.805 | 0.480 | 24.337 | 45.301 | 94.123 |
| ano.34.LR | 44.416 | 0.471 | 22.774 | 43.638 | 89.277 |

Table S2. Frequencies with which the probability of the chi-squared estimate of deviance change was the lowest, and the delta-AIC value was the highest, between compared models.

| | MP1 | | | MP2 | | |
|---|---|---|---|---|---|---|
| | ano.p | dAIC | LR | Ano.p | dAIC | LR |
| 12 | 1 | 0 | 0 | 0 | 0 | 0 |
| 13 | 6 | 3 | 0 | 0 | 0 | 0 |
| 14 | 455 | 475 | 500 | 5 | 13 | 500 |
| 23 | 0 | 0 | 0 | 0 | 0 | 0 |
| 24 | 20 | 14 | 0 | 57 | 114 | 0 |
| 34 | 18 | 8 | 0 | 438 | 373 | 0 |

### Appendix S4. Liana densities at the species level

Table S1. Mean and SE of number of lianas per tree (≥ 30 cm gbh) at Danum in 1988 and 2018, for those species with $n \geq 20$ trees per species in the two plots (MP1 and MP2) combined. Nomenclature follows the revisions of 2015. Abbreviations (Abbr.) are used in Figs 4-6 in the main paper, and Fig. S1 in this appendix.

| Species name | Authority | Family | Abbr. | 1988 | | | 2018 | | |
|---|---|---|---|---|---|---|---|---|---|
| | | | | *n* | mean | se | *n* | mean | se |
| *Aglaia silvestris* | (M.Roem.) Merr. | Meliaceae | Asi | 20 | 2.20 | 0.91 | | | |
| *Aporosa falcifera* | Hook.f. | Phyllanthaceae | Af | 122 | 2.90 | 0.31 | 85 | 1.84 | 0.17 |
| *Ardisia sanguinolenta* | Blume | Primulaceae | Asa | 30 | 1.33 | 0.35 | 38 | 1.37 | 0.23 |
| *Baccaurea tetrandra* | (Baill.) Müll.Arg. | Phyllanthaceae | Bt | 77 | 2.42 | 0.32 | 79 | 2.24 | 0.23 |
| *Barringtonia lanceolata* | (Ridl.) Payens | Lecythidaceae | Bl | 76 | 2.00 | 0.32 | 66 | 1.61 | 0.22 |
| *Canarium denticulatum* | Blume | Burseraceae | Ca | | | | 20 | 2.20 | 0.57 |
| *Chisocheton sarawakanus* | (C.DC.) Harms | Meliaceae | Cs | 51 | 3.39 | 0.59 | 53 | 1.98 | 0.31 |
| *Cleistanthus contractus* | Airy Shaw | Phyllanthaceae | Cc | | | | 20 | 0.85 | 0.27 |
| *Dacryodes rostrata* | (Blume) H.J.Lam | Burseraceae | Dr | 27 | 3.07 | 0.56 | 26 | 1.73 | 0.41 |
| *Dimorphocalyx muricatus* | (Hook.f.) Airy Shaw | Euphorbiaceae | Dm | 62 | 1.87 | 0.34 | 76 | 1.14 | 0.15 |
| *Diospyros elliptifolia* | Merr. | Ebenaceae | De | 20 | 1.60 | 0.42 | | | |
| *Dipterocarpus kerrii* | King | Dipterocarpaceae | Dk | 25 | 1.32 | 0.64 | 25 | 0.52 | 0.16 |
| *Drypetes longifolia* | (Blume) Pax & K.Hoffm. | Putranjivaceae | Dl | 32 | 1.72 | 0.50 | | | |
| *Dysoxylum cyrtobotryum* | Miq. | Meliaceae | Dc | 49 | 1.57 | 0.40 | 82 | 1.90 | 0.26 |
| *Dysoxylum rigidum* | (Ridl.) Mabb. | Meliaceae | Do | | | | 20 | 1.60 | 0.39 |
| *Fordia splendidissima* | (Miq.) Buijsen | Leguminosae | Fs | | | | 24 | 1.46 | 0.36 |
| *Gironniera nervosa* | Planch. | Cannabaceae | Gn | | | | 39 | 1.51 | 0.42 |
| *Gonystylus keithii* | Airy Shaw | Thymelaeaceae | Gk | 31 | 2.48 | 0.59 | 36 | 2.56 | 0.38 |
| *Hancea penangensis* | (Müll.Arg.) Sierra et al. | Euphorbiaceae | Hpe | 36 | 1.97 | 0.43 | 34 | 1.32 | 0.25 |
| *Hancea stipularis* | (Airy Shaw) Sierra et al. | Euphorbiaceae | Hs | 30 | 1.70 | 0.49 | 21 | 2.10 | 0.54 |

| | | | | | | | | | |
|---|---|---|---|---|---|---|---|---|---|
| *Hopea nervosa* | King | Dipterocarpaceae | Hn | | | | 33 | 1.55 | 0.43 |
| *Hydnocarpus polypetalus* | (Slooten) Sleumer | Flacourtiaceae | Hpo | | | | 24 | 1.42 | 0.30 |
| *Knema latericia* | Elmer | Myristicaceae | Kla | | | | 21 | 1.90 | 0.36 |
| *Koilodepas laevigatum* | Airy Shaw | Euphorbiaceae | Kle | 29 | 0.83 | 0.30 | 24 | 0.83 | 0.22 |
| *Lithocarpus gracilis* | (Korth.) Soepadmo | Fagaceae | Lgr | 27 | 2.15 | 0.71 | | | |
| *Lithocarpus leptogyne* | (Korth.) Soepadmo | Fagaceae | Ll | 28 | 2.46 | 1.07 | 29 | 1.45 | 0.31 |
| *Lithocarpus nieuwenhuisii* | (Seemen) A.Camus | Fagaceae | Ln | 57 | 2.46 | 0.56 | 49 | 1.86 | 0.38 |
| *Litsea caulocarpa* | Merr. | Lauraceae | Lc | 39 | 1.64 | 0.33 | 54 | 1.50 | 0.23 |
| *Litsea garciae* | Vidal | Lauraceae | Lga | | | | 23 | 2.09 | 0.47 |
| *Litsea ochracea* | (Blume) Boerl. | Lauraceae | Lo | 36 | 2.08 | 0.48 | 20 | 1.15 | 0.35 |
| *Lophopetalum beccarianum* | Pierre | Celastraceae | Lb | 20 | 1.75 | 0.61 | 58 | 0.81 | 0.14 |
| *Maasia sumatrana* | (Miq.) Mols et al. | Annonaceae | Ms | 79 | 2.58 | 0.34 | 70 | 2.04 | 0.27 |
| *Madhuca korthalsii* | (Pierre ex Burck) H.J.Lam | Sapotaceae | Mk | 115 | 2.88 | 0.32 | 126 | 2.17 | 0.18 |
| *Mallotus wrayi* | King ex Hook.f. | Euphorbiaceae | Mw | 202 | 2.33 | 0.20 | 207 | 1.60 | 0.13 |
| *Microcos reticulata* | Ridl. | Malvaceae | Mr | 30 | 1.33 | 0.40 | 31 | 1.48 | 0.31 |
| *Neoscortechinia philippinensis* | (Merr.) Welzen | Euphorbiaceae | Np | 58 | 1.41 | 0.38 | 35 | 1.57 | 0.48 |
| *Nothaphoebe species a* | --- | Lauraceae | Nsp | 20 | 1.20 | 0.44 | | | |
| *Ochanostachys amentacea* | Mast. | Olacaceae | Oa | 30 | 1.53 | 0.48 | 26 | 1.38 | 0.35 |
| *Parashorea malaanonan* | Merr. | Dipterocarpaceae | Pm | 66 | 1.42 | 0.33 | 59 | 0.64 | 0.16 |
| *Pentace laxiflora* | Merr. | Malvaceae | Pl | 95 | 2.53 | 0.40 | 96 | 1.68 | 0.18 |
| *Polyalthia cauliflora* | Hook.f. & Thomson | Annonaceae | Pca | 39 | 1.18 | 0.31 | 23 | 0.96 | 0.27 |
| *Polyalthia congesta* | (Ridl.) J.Sinclair | Annonaceae | Pco | 21 | 2.10 | 0.66 | 21 | 1.10 | 0.24 |
| *Polyalthia rumphii* | (Blume ex Hensch.) Merr. | Annonaceae | Pr | 21 | 1.14 | 0.31 | 45 | 1.24 | 0.31 |
| *Polyalthia xanthopetala* | Merr. | Annonaceae | Px | 29 | 1.83 | 0.60 | | | |
| *Pternandra galeata* | Ridl. | Melastomataceae | Pg | 22 | 1.73 | 0.50 | | | |
| *Reinwardtiodendron humile* | (Hassk.) Mabb. | Meliaceae | Rh | 31 | 1.97 | 0.34 | | | |
| *Scorodocarpus borneensis* | (Baill.) Becc. | Olacaceae | Sb | 55 | 2.93 | 0.53 | 58 | 1.72 | 0.21 |
| *Shorea argentifolia* | Symington | Dipterocarpaceae | Sa | 28 | 1.57 | 0.68 | 29 | 0.69 | 0.38 |
| *Shorea fallax* | Meijer | Dipterocarpaceae | Sf | 75 | 1.71 | 0.42 | 86 | 1.06 | 0.15 |
| *Shorea johorensis* | Foxw. | Dipterocarpaceae | Sj | 76 | 1.43 | 0.38 | 75 | 0.75 | 0.18 |
| *Shorea leprosula* | Miq. | Dipterocarpaceae | Sl | 29 | 0.97 | 0.37 | 25 | 0.36 | 0.18 |
| *Shorea parvifolia* | Dyer | Dipterocarpaceae | Spr | 106 | 1.42 | 0.34 | 115 | 0.63 | 0.15 |

| | | | | | | | | | |
|---|---|---|---|---|---|---|---|---|---|
| *Shorea pauciflora* | King | Dipterocarpaceae | Spu | 33 | 2.85 | 0.58 | 31 | 1.84 | 0.35 |
| *Shorea pilosa* | P.S.Ashton | Dipterocarpaceae | Spi | 47 | 0.94 | 0.32 | 57 | 0.72 | 0.18 |
| *Syzygium castaneum* | (Merr.) Merr. & L.M.Perry | Myrtaceae | Sc | | | | 20 | 1.90 | 0.51 |
| *Syzygium elopurae* | (Ridl.) Merr. & L.M.Perry | Myrtaceae | Se | 30 | 1.80 | 0.46 | | | |
| Syzygium kunstleri | (King) Bahadur & R.C.Gaur | Myrtaceae | Sk | 33 | 3.39 | 0.76 | 29 | 1.97 | 0.34 |
| Syzygium lineatum | (DC.) Merr. & L.M.Perry | Myrtaceae | Sl | 31 | 1.68 | 0.43 | 30 | 2.23 | 0.39 |
| Syzygium peregrinum | (Blume) Merr. & L.M.Perry | Myrtaceae | Sp | 48 | 3.10 | 0.57 | 39 | 1.54 | 0.35 |
| Vatica dulitensis | Symington | Dipterocarpaceae | Vd | 26 | 2.00 | 0.76 | 21 | 1.29 | 0.34 |
| Xanthophyllum vitellinum | (Blume) D.Dietr. | Polygalaceae | Xv | 25 | 2.32 | 0.55 | 30 | 1.63 | 0.33 |

Fig. S1. Relationships between mean species' number of lianas per tree (nr_lianas) and the stem relative growth rate (rgr, mm/m/yr), for species with ≥ 20 trees in plots MP1 and MP2 combined in (a) 1988 and (b) 2018. Closed red circles, Dipterocarpaceae; open blue squares, Euphorbiaceae; open green triangles, other families. This figure complements Fig. 4 of the main text.

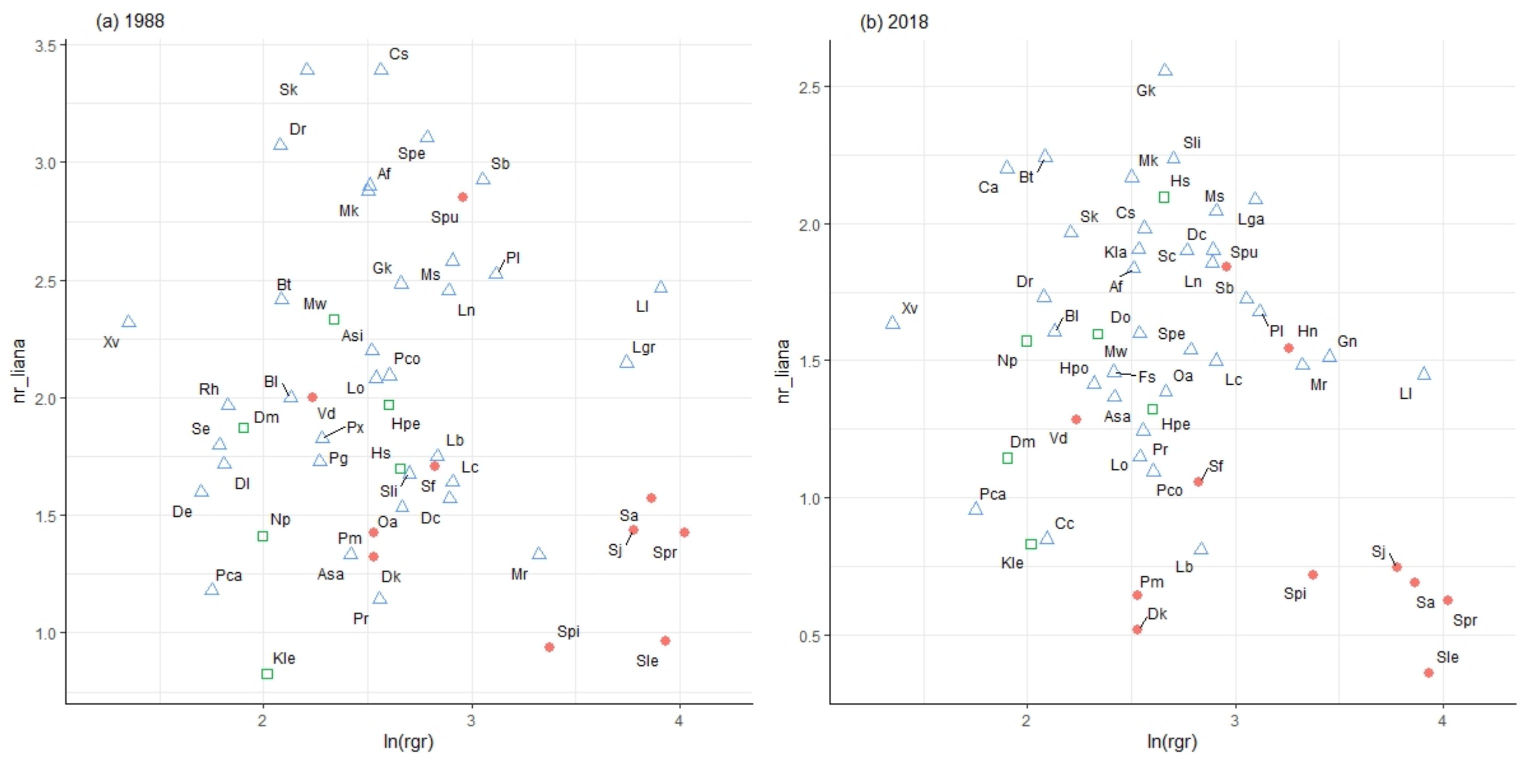

Ecosphere/Changes in liana density over 30 years in a Bornean rain forest supports the escape hypothesis/D. M. Newbery and C. Zahnd

**Appendix S5. Code for statistical analyses**

1. R-code for the randomization functions (written by CZ)

```
# Change in liana abundance with time. A and B are auxiliary functions used later in C-E.
# In C-E extraction of parameter estimates and statistics was omitted, in E the model
# comparisons with ANOVA were also omitted. C-E were run for each plot, and for
# dipterocarps and euphorbs in each plot separately.

# A. Randomly sample close to half of the rows (tag is the tree numbers).
randomRows <- function(dat, n = round(((1/2) * nrow(dat)) -1, 0)){
	sample(dat$tag, n, replace = F)
} # end of randomRows function.

# B. Create a flagging column in the dataset based on randomRows (A).
flagging <- function(dat, f){
for(i in 1:nrow(dat)){
if(dat$tag[i] %in% f){
dat$flag[i] <- 1
}else{
dat$flag[i] <- 0
}
}
return(dat)
} # end of flagging function.

# C. Change in proportion of liana infested trees (df is the input data frame)
randomProp <- function(df, iter = 500){
set.seed(8751)
# initialize output vectors, e.g.:
p_val <- rep(NA, iter)
# Run the 500 random subsamplings:
for(i in 1:iter){
# randomly select close to half of all rows and create a flagging column:
temp <- randomRows(dat = df)
flagged <- flagging(dat = df, f = temp)
# subset data based on flagging column for 1988 and 2018:
flagged88 <- subset(flagged, flagged$flag == 1)
flagged88 <- subset(flagged88, !is.na(flagged88$liana_pres88))
flagged18 <- subset(flagged, flagged$flag == 0)
flagged18 <- subset(flagged18, !is.na(flagged18$liana_pres18))
# Calculate total number of trees and proportions with lianas (liana_pres indicate liana
# presence (1) or absence (0) on each tree):
proportions <- integer(2)
```

```
proportions[2] <- sum(flagged88$liana_pres88)
proportions[1] <- sum(flagged18$liana_pres18)
lengths <- integer(2)
lengths[2] <- nrow(flagged88)
lengths[1] <- nrow(flagged18)
# The chi-squared test:
test <- prop.test(proportions, lengths, correct = TRUE)
# save results into output vectors, e.g.:
p_val[i] <- round(test$p.value, 6)
}
# here all output vectors can be put together into a data.frame and returned from the
# function.
} # end of randomProp function.
```

D. Frequency of trees in liana density classes.

```
randomCount <- function(df, iter = 500){
set.seed(8751)
# initialize output vectors, e.g.:
p_val <- rep(NA, iter)
# randomly select close to half of all rows and create a flagging column:
for(i in 1:iter){
temp <- randomRows(dat = df)
flagged <- flagging(dat = df, f = temp)
# subset data based on flagging column for 1988 and 2018:
flagged88 <- subset(flagged, flagged$flag == 1)
flagged88 <- subset(flagged88, !is.na(flagged88$nr_liana88))
flagged18 <- subset(flagged, flagged$flag == 0)
flagged18 <- subset(flagged18, !is.na(flagged18$nr_liana18))
# Find the most liana count classes possible with no class having < 5 counts in either year.
maxi88 <- max(flagged88$nr_liana88)
maxi18 <- max(flagged18$nr_liana18)
maxi <- integer(1)
maxi <- max(c(maxi88, maxi18))
classes <- c(0:maxi)
counts88 <- integer(length(classes))
for(j in 1:(length(classes))){
counts88[j] <- nrow(flagged88[flagged88$nr_liana88 == (j-1),])
}
counts18 <- integer(length(classes))
for(k in 1:(length(classes))){
counts18[k] <- nrow(flagged18[flagged18$nr_liana18 == (k-1),])
}
cs <- factor(0:maxi)
s <- 1
while(counts88[s] >= 5){
s <- s + 1
}
```

```
t <- 1
while(counts18[t] >= 5){
t <- t + 1
}
shorten <- min(c(s, t))
}
cs[shorten:length(classes)] <- as.character(cs[shorten])
# Sum the trees in the liana density classes defined above:
of88 <- as.vector(tapply(counts88, cs, sum))
of18 <- as.vector(tapply(counts18, cs, sum))
of <- c(of88, of18)
mtrx <- matrix(of, nrow = length(of88))
mtrx <- mtrx[1:shorten,]
# run the chi-squared test:
test <- chisq.test(mtrx)
# save results into output vectors, e.g.:
p_val[i] <- test$p.value
}
# here all output vectors can be put together into a data.frame and returned from the
# function.
} # end of randomCount function.

# E. Mean number of lianas per tree:
randomMean <- function(df, iter = 500){
set.seed(8751)
# initialize output vectors for all 4 models and model comparisons with ANOVA, e.g.:
inter.zval.1 <- vector(mode="numeric",iter)
year.se.2 <- vector(mode="numeric",iter)
# randomly select close to half of all rows and create a flagging column:
for(i in 1:iter){
temp <- randomRows(dat = df)
flagged <- flagging(dat = df, f = temp)
# subset data based on flagging column for 1988 and 2018 and recombine subset data:
flagged88 <- subset(flagged, flagged$flag == 1)
flagged88 <- subset(flagged88, !is.na(flagged88$nr_liana88))
flagged88a <- subset(flagged88, select = - c(size_class18, liana_pres18, bamboo18,
                                nr_liana18, stat18, GBH18, bamboo88))
   names(flagged88a) <- c("plot", "subplot", "tag", "f_code15" ,"g_code15",
                  "sp_code15", "GBH", "stat", "nr_liana", "uni_code",
                  "liana_pres", "size_class", "flag")
flagged88a$year <- as.factor("88")
flagged18 <- subset(flagged, flagged$flag == 0)
flagged18 <- subset(flagged18, !is.na(flagged18$nr_liana18))
flagged18a <- subset(flagged18, select = - c(size_class88, liana_pres88, bamboo18,
                                nr_liana88, stat88, GBH88, bamboo88))
   names(flagged18a) <- c("plot", "subplot", "tag", "f_code15", "g_code15",
                  "sp_code15", "uni_code", "GBH", "stat", "nr_liana",
```

```
                    "liana_pres", "size_class", "flag")
flagged18a$year <- as.factor("18")
comb <- rbind(flagged88a, flagged18a)
# create lnGBH:
comb$lnGBH <- log(comb$GBH)
comb$lnGBHq <- (comb$lnGBH)^2
# run the 4 glm.nb models:
mod1 <- with(comb, glm.nb(nr_liana ~ year, link = log))
mod2 <- with(comb, glm.nb(nr_liana ~ year + lnGBH, link = log))
mod3 <- with(comb, glm.nb(nr_liana ~ year * lnGBH, link = log))
mod4 <- with(comb, glm.nb(nr_liana ~ year * lnGBH + lnGBHq + year:lnGBHq, link = log))
# calculate pseudo R-squared for each model, e.g.:
LLM <- -(summary(mod1)$deviance)/2
LL0 <- -( summary(mod1)$null.deviance)/2
pseudoR2.1 <- ((LL0-LLM)/LL0)*100
rm(“LLM”, “LL0”) # (do the same for the other models)
# here pairwise model comparisons with anova would be made as described in the main
# text.
# here all parameter estimates and statistics from the models and model comparisons
# would be saved in the output vectors.
}
# here all output vectors can be put together into a data.frame and returned from the
# function.
} # end of randomMean function.
```

### 2. Regression modelling with spatial autocorrelation

Essential R-commands (applied by DMN).

```
# basic input/variable calcs and summary/anova/plotting/etc commands not shown.

# 'liana2er.88' or 'liana2er.18' are base data.frames,'liana2er.88s' or 'liana2er.18s'

# those for survivors. 'elevplus' has within-plot elevations of 1988 added to 2018 trees.


# 1. nr_lianas (variable names for 1988 -- similar for 2018). [GBH is girth at breast height]

# calculate distance matrix (plot/census specific)

distMat <- as.matrix(dist(cds,method="euclidean",diag=TRUE,upper=TRUE,p=2))

distMat.inv <- ifelse(distMat == 0, 0, 1/distMat)

# negative binomial glm (run for census x plot separately)[lnGBH88q is lnGBH88 squared]

nb_mod.88 <-
glm.nb(formula=nr_liana88~lnGBH88+lnGBH88q+elevplus,data=liana2er.88,link=log)

# hglm with spatial correlation matrix (run for census x plot separately)

hglm_mod.88 <- hglm(fixed = nr_liana18~lnGBH88+lnGBH88q+elevplus,

 rand.family = SAR(D = distMat.inv),

 data = liana2er.88,random = ~1|tag,family = quasipoisson(link="log"))

# glmmML (for family level analyses): spatial neighbours as subplot coords (20-m x 20-m grids).

mod1s <- glmmML(surv18~nr_liana.sqrt+lnGBH88+elevplus,cluster = subpl, family=binomial)


# 2. tree survival (nr_liana.sqrt is square-root of number of lianas)

# binomial glm (run for plots separately)[surv18 is survival 1988-2018]

bin_mod <- glm(surv18~nr_liana.sqrt+lnGBH88+elevplus,binomial)

# autologistic regression with distance-based correlation term

autol.mod <- logistic.regression(ldata=liana2er.88,y='surv18',x=c('nr_liana.sqrt',

 'lnGBH88','elevplus'), penalty=TRUE,autologistic=TRUE,type="inverse.squared",

 coords=cds,bw=20,style="B")
```

```
# 3. tree growth (run for plots separately)[rgr18 is relative growth rate 1988-2018]
# gaussian gls, or equvalently lm, regression
gls.mod <- gls(rgr18~nr_liana.sqrt+lnGBH88,liana2er.88s)
# A. gls regression with variogram-based spatial correlation function
gls_mod.exp <- update (gls.mod, corr = corExp(c(60,0.9), form = ~ x + y, nugget=T))
# B. spatial lm regression (using distance to nearest neighbours, defined by Moran's I
# and Geary's C statistics.
cds.nb <- dnearneigh(as.matrix(cds),d1=0,d2=20)
cds.lw <- nb2listw(cds.nb,style="B",zero.policy=TRUE)
spatlm.mod <- spautolm(rgr18 ~ nr_liana.sqrt + lnGBH88, data = liana2er.88s,
 family = "SAR", listw = cds.lw, zero.policy = TRUE)
```
